# mini-Complexome Profiling (mCP), an FDR-controlled workflow for global targeted detection of protein complexes

**DOI:** 10.1101/2024.04.22.590599

**Authors:** Hugo Amedei, Niels Paul, Brian Foo, Lisa Neuenroth, Stephan E. Lehnart, Henning Urlaub, Christof Lenz

## Abstract

**Introduction:** Co-fractionation mass spectrometry couples native-like separations of protein/protein complexes with mass spectrometric proteome analysis for global characterization of protein networks. The technique allows for both de novo detection of complexes and for the detection of subtle changes in their protein composition. The typical requirement for fine-grained fractionation of >80 fractions, however, translates into significant demands on sample quantity and mass spectrometric instrument time, and represents a significant barrier to experimental replication and the use of scarce sample material (ex. Patient biopsies).

**Methods:** We developed mini-Complexome Profiling (mCP), a streamlined workflow with reduced requirements for fractionation and, thus, biological material and laboratory and instrument time. Soluble and membrane-associated protein complexes are extracted from biological material under mild conditions, and fractionated by Blue Native electrophoresis using commercial equipment. Each fraction is analyzed by data independent acquisition mass-spectrometry, and known protein complexes are detected based on the coelution of known components using a novel R package with a controlled false discovery rate approach. The tool is available to the community on a GitHub repository.

**Results:** mCP was benchmarked using HEK293 cell lysate and exhibited performance similar to established workflows, but from a significantly reduced number of fractions. We then challenged mCP by performing comparative complexome analysis of cardiomyocytes isolated from different chambers from a single mouse heart, where we identified subtle chamber-specific changes in mitochondrial OxPhos complexes.

**Discussion:** The reduced sample and instrument time requirements open up new applications of co-fractionation mass spectrometry, specifically for the analysis of sparse samples such as human patient biopsies. The ability to identify subtle changes between similar tissue types (left/right ventricular and atrial cardiomyocytes) serves as a proof of principle for comparative analysis of mild/asymptomatic disease states.

## 1 Introduction

On the molecular level, the majority of cellular functions such as energy metabolism, cell division or replication rely on the actions of protein-protein complexes (PPCs). The systematic study of PPCs is therefore crucial to understanding and characterizing cellular biology. Over the last decade, complexome profiling (CP) as an implementation of co-fractionation mass spectrometry (CF-MS) has emerged as a key approach to detecting PPCs on a global scale. CP frequently includes mild non-ionic detergent-based solubilization and extraction of complexes; non-denaturing (‘native-like’) enrichment and fractionation of the resulting PPCs; profiling of the protein contents across all fractions by mass spectrometry; and finally, statistical and bioinformatic analysis of the fraction-resolved protein profiles obtained. The co-elution of proteins suggests their biophysical interaction in a non-covalent PPC, and there are computational approaches to calculate co-elution scores and correlate them to protein-protein interaction (Bludau et al., 2020; Giese et al., 2015; Havugimana et al., 2012; Heusel et al., 2019; Stacey et al., 2017). CP has been used to complement affinity purification experiments (Uliana et al., 2023) and is currently the method of choice for proteome-wide screening of interactome rewiring across samples or conditions (Bludau, 2021). Recently, efforts have begun to consolidate this young technique to enable better sharing of biological results, which will be of great value to the scientific community (Van Strien et al., 2021).

All complexome profiling methods are based on the premise of successful solubilization of intact PPCs from their biological environments with high fidelity. This is particularly challenging in the case of membrane-associated PPCs, where a 3-step extraction mechanism has been proposed (Le Maire et al., 2000): in stage I, detergent molecules interact with membrane lipids, and non-micellar detergent partitions into the phospholipid bilayer are observed. In stage II, a mixture of phospholipid-detergent micelles and phospholipid membranes saturated with detergent coexist in equilibrium. Finally, in stage III, membrane complexes are fully solubilized into micelles. Therefore, membrane-associated PPCs may or may not be characterized by a defined stoichiometry. When using ‘native-like’ separations for their enrichment, this often results in poorly defined elution profiles compared to those observed for soluble complexes, making it challenging to establish statistical parameters for their automated detection.

The study of membrane-associated PPCs has gained particular attention in the field of cardiovascular research, where they play key roles in several key functions such as Ca^2+^ homeostasis, cell-cell junctions and energy metabolism. A frequently used approach involves the preparation of so-called enriched membrane fractions (Santos et al., 2002; Soni et al., 2016), followed by solubilization with non-ionic detergents such as cholate, deoxycholate, 3-((3-cholamidopropyl)dimethylamino)-1-propane sulfonate (CHAPS) or mixtures thereof (Cornelius, 1991). Alternatively, enriched membrane fractions may be solubilized by glycosides such as digitonin, a strategy frequently used for the detection of mitochondrial (Schtigger et al., 1991) or cardiac membrane proteins (Alsina et al., 2019). E.g., mitochondrial supercomplexes were visualized using blue native electrophoresis (BNE) following mild digitonin or Nonoxinol 40 (NP-40) extraction (Acín-Pérez et al., 2008), or using size exclusion chromatography (SEC) followed by octaethylene glycol monododecyl ether (C12E8) extraction (Foo et al., 2024). In the context of cardiac research, the compositions of active mitochondrial human respiratory super-complexes composed of mitochondrial complex I, III, and IV (CI1III2IV1) and the architecture of mitochondrial complex I, II, and IV(CI2III2IV2) were elucidated (Guo et al., 2017). This is of particular importance since the metabolic state of e.g. mouse hearts has been directly linked to the abundance of mitochondrial supercomplexes in isolated mitochondria (Zheng et al., 2023). As an alternative to the preparation of enriched membrane fractions, cytoplasmic and membrane-associated protein complexes can be extracted directly from whole cells. In this case, the lysis buffer is applied directly to the cells without previous fractionation, an approach here called general lysis (Heusel et al., 2019). It is worth mentioning that both enriched membrane fractions and general lysis methods are able to extract both membrane and soluble protein complexes, albeit with varying efficiencies and fidelities.

Once complexes are solubilized, the fractionation of PPCs is carried out using non-denaturing or ‘native-like’ separation methods such as BNE (Alsina et al., 2019; Heide et al., 2012), SEC (Heusel et al., 2019; Heusel et al., 2020), ion exchange chromatography (IEX) (Havugimana et al., 2012) or density gradient centrifugation (Páleníková et al., 2021). Among these, BNE and SEC are by far the most frequently used due to their demonstrated potential for high proteome and PPC coverage (Skinnider and Foster, 2021). Blue Native polyacrylamide gel electrophoresis (BN-PAGE) was originally developed as a variant of native PAGE to study mitochondrial protein complexes. Depending on the gel matrix used, the method enables separations up to the 10 MDa molecular weight range (Schtigger et al., 1991; Wittig et al., 2006). Solubilizing detergents are not strictly required for BN-PAGE; the combination of the anionic Coomassie dye with non-ionic detergents, however, can generate an amphipathic effect that can be detrimental to protein complex stability (Wittig et al., 2006), necessitating careful optimization.

Historically, mass spectrometric analysis in CP-MS experiments has mostly been performed using data-dependent acquisition mass spectrometry (DDA-MS) (Skinnider and Foster, 2021). In the last decade, however, data-independent acquisition mass spectrometry (DIA-MS) has emerged as a superior acquisition mode in proteomics to detect and quantify proteins in complex samples (Gillet et al., 2012). Due to its sensitivity (Bruderer et al., 2015; Dowell et al., 2021)] and quantitative fidelity (Barkovits et al., 2020), DIA-MS has been increasingly applied to complexome profiling (Bludau et al., 2020; Heusel et al., 2020 and 2019). More recently, trapped ion mobility-time of flight-mass spectrometry (timsToF-MS) (Meier et al., 2015) has expanded the utility of DIA-MS by introducing an additional dimension of separation to the analysis of peptide precursors. The approach, termed diaPASEF (Meier et al., 2020), allows for the identification and quantification of high numbers of proteins in very short liquid chromatography tandem mass spectrometry (LC/MS/MS) experiments gradients (Skowronek et al., 2022), and can be tailored to different gradient lengths and sample complexities and novel DIA setup designs (Szyrwiel et al., 2022). Following the development and refinement of these acquisition paradigms, multiple data analysis platforms have been made available for the analysis of DIA data, including DIA-NN (Demichev et al., 2020), Spectronaut (Bruderer et al., 2015), Skyline (MacLean et al., 2010), Max Quant (Sinitcyn et al., 2021) and MS-Fragger (Yu et al., 2022).

The statistical and bioinformatic analysis of CF-MS datasets has been greatly improved, and today there are many different alternatives to detect PPCs in complex biological samples. The detection of protein complexes by unbiased clustering of protein abundance profiles, known as non-targeted analysis, is implemented in software tools like NOVA (Giese et al., 2015). In addition, there have been approaches to detect protein complexes in LC/MS/MS that look for protein complexes by machine learning, namely EPIC (Hu et al., 2019), PCprophet (Fossati et al., 2021b), ComplexFinder (Nolte and Langer, 2021), PrInCE (Stacey et al., 2017), and DIP-MS (Frommelt et al., 2023). Targeted analysis, on the other hand, relies on the introduction of a working hypothesis or ‘ground truth’ to extract protein profiles for further correlation analysis (Bludau, 2021). This principle is implemented in software packages like CCprofiler (Bludau et al., 2020) and Complexbrowser (Michalak et al., 2019). Ground truth is often introduced from existing PPC databases such as CORUM (Tsitsiridis et al., 2023), String (Szklarczyk et al., 2023) or Complex Portal (Meldal et al., 2015).

The main current disadvantage of most CP approaches is the classic requirement for high numbers of fractions, which in turn requires significant mass spectrometric instrument time for analysis. Most established approaches use 50-80 fractions, and high-resolution experiments with up to 1,000 fractions have been tested (Havugimana et al., 2012). The resulting requirement for MS instrument time has been greatly reduced due to the advent of high density, short gradient LC/MS/MS methods such as diaPASEF, and systematic studies have demonstrated the limited benefits of over-fractionation, as pointed out in a meta-study of 206 published datasets (Skinnider and Foster, 2021). Even so, the effort required currently precludes the comparison of significant numbers of samples, or even replication to assess the reproducibility of quantitation.

Here, we introduce a novel workflow called mini-Complexome Profiling (mCP), complete with a publicly available R package available for data analysis, which allows for targeted detection of known protein complexes in low fractionation, BNE-based experiments. At 35 fractions and correspondingly reduced amounts of required instrument time, mCP is particularly suitable for screening samples from multiple conditions, and allows for replication to stabilize results. We developed and benchmarked it on HEK293 cells, an established model for protein complex studies, where our approach allowed the detection of 535 protein complexes in total cell lysates, equivalent to the number of PPCs detected in 81 fractions previously (Bludau et al., 2020; Heusel et al., 2019). Subsequently, we show the versatility of mCP in the comparative analysis of cardiomyocytes from the left and right ventricles as well as the atria of a single mouse heart, demonstrating its utility in performing comparative analysis from sparse samples.

## 2 Materials and Methods

### 2.1 HEK293 Cultivation

HEK293A cells (ATCC) were cultivated on T75 flasks (Greiner) in low-glucose Dulbecco’s Modified Eagle’s Medium (DMEM, Sigma) supplemented with 10% (v/v) fetal bovine serum (FBS, Thermo Fisher), 2 mM L-glutamine GlutaMAX (Sigma-Aldrich) and 1x penicillin/streptomycin (Sigma-Aldrich). Cells were cultivated under humidified conditions with 5% CO2 at 37°C and passaged every 3-4 days using 0.05% trypsin/EDTA (Sigma-Aldrich). Cells were harvested in ice-cold PBS buffer using a cell scraper, and sedimented for 5 min at 500x G at 4°C.

### 2.2 Murine Cardiomyocyte Isolation

Animal experiments were performed in accordance with directive 2010/63/EU of the European Parliament and in keeping with NIH guidelines. The use of WT mice as a source of tissues for assay development is covered by the institute’s animal protocols 11/2 and 22/19, reviewed and approved by the institutional animal committee of the University Medical Center Göttingen, and finally approved by the veterinarian state authority (LAVES, Oldenburg, Germany).

A 12-week old, male, wildtype mouse with a C57BL6N genotype (ID: OG08-1244, line PLB-KO) was sacrificed and cardiomyocytes were isolated using a published protocol (Wagner et al., 2014). In brief, the proximal aorta was cannulated using a 21-gauge needle and connected to a modified Langendorff perfusion setup. Hearts were perfused at a constant flow of 4 ml/min with a nominally Ca^2+^-free perfusion buffer for 4 min at 37°C, followed by collagenase containing buffer for another 9 min at 37°C. The left and right ventricles as well as the atria were dissected under a binocular microscope (Stemi 305, Zeiss) in 2 ml digestion buffer, and digestion was stopped by adding 3 ml stopping buffer containing 10% FBS. Isolated cardiomyocytes from the left and right ventricle (LVCM/RVCM) and atria (ACM) were washed twice with the stopping buffer, cells sedimented for 8 min by gravity at room temperature (RT), the supernatant was discarded and the cell pellets were stored at -80°C.

### 2.3 Non-denaturing Lysis and BNE Fractionation

Protein complexes were extracted from HEK293 cells and isolated cardiomyocyte samples following established protocols (Bludau et al., 2020), with the omission of the concentration and buffer exchange steps. Lysis was performed at 4°C until the electrophoresis step. HEK293 cells were resuspended in 500 μl of HNN lysis buffer (0.05% vol/vol NP-40, 50 mM HEPES, 150 mM NaCl, 50 mM NaF, 200 μM Na_3_VO_4_, 1 mM phenylmethylsulfonyl fluoride (PMSF), and 1x protease inhibitor cocktail, pH 7.5) (Bludau et al., 2020). Cardiomyocytes isolated from the mouse left and right ventricle and from the atria were lysed by addition of 0.62 μl of non-denaturing lysis buffer per mg of moist pellet. Lysates were cleared by ultracentrifugation (100,000x G, 15 min) and protein concentrations were determined by BCA (Pierce) and determined to be at least 2.7 mg/ml. Lysates were analyzed by Blue Native-PAGE on pre-cast Native 3-12% Bis-Tris minigels (Invitrogen) using standard protocols (Wittig et al., 2006). Nativemark standard (Invitrogen) was used for molecular weight calibration. 35 μg of protein per lane were used for HEK293 complexome profiling, and 60 μg for cardiomyocytes. Lanes were cut into 35 equidistant fractions irrespective of staining using an in-house stainless steel cutter. All fractions were reduced, alkylated, and digested with trypsin in 96 well microplates as previously described (Atanassov and Urlaub, 2013).

### 2.4 SDS-PAGE Purification

Briefly, 17.6 μg cleared lysate from HEK293 cells, or 5 μg cleared lysate from cardiomyocytes were purified by a brief 1 cm SDS-PAGE run (Nu-PAGE minigel 4-12%, Invitrogen). After Coomassie staining, each lane was excised in one piece, reduced, alkylated, and digested with trypsin as previously described (Atanassov and Urlaub, 2013).

### 2.5 LC/MS/MS Data Acquisition

In-gel digested samples were resuspended via sonication in loading buffer (2% aqueous acetonitrile, 0.1% formic acid) and spiked with iRT standard peptides (1/100 diluted, Biognosys). Peptides were separated on a nanoflow chromatography system using reversed-phase chromatography, and the eluent was analyzed on a hyphenated timed ion mobility-time of light mass spectrometer (timsTOF Pro 2, Bruker) in either PASEF or diaPASEF acquisition modes.

#### 2.5.1 Data-Independent Acquisition for Proteome Profiling

Peptides were separated using a linear 12.5 min gradient of 4-32% aqueous acetonitrile versus 0.1% formic acid on a reversed phase-C18 column (PepSep Fifteen, 150x0.150 mm, Bruker) at a flow rate of 850 nl min^-1^. DIA analysis was performed in diaPASEF mode using a customized 8x2 window acquisition method from *m/z* 100 to 1700, and from 1/K_0_=0.7-1.5 to include the 2+/3+/4+ population in the *m/z*-ion mobility plane (Skowronek et al., 2022). The collision energy was ramped linearly as a function of the mobility from 59 eV at 1/K_0_=1.5 Vs cm^−2^ to 20 eV at 1/K_0_=0.7 Vs cm^−2^. Two to four technical replicates of each sample were acquired.

#### 2.5.2 Data-Independent Acquisition for Spectral Library Generation

Peptides coming from lysates were measured using two different gradients and columns. The first two technical replicates of each lysate were acquired with the same column and gradient as DIA for proteome profiling (see above). Another two technical replicates were measured using a longer linear 100 min gradient of 2-37% aqueous acetonitrile versus 0.1% formic acid on a reversed phase-C18 column (Aurora Elite 250x0.075 mm, IonOpticks) at a flow rate of 250 nl min^-1^ and the same DIA setting used for proteome profiling.

### 2.6 Data Processing

DIA complexome profiling datasets were analyzed by DIA-NN software v1.8.1 (Charité; Demichev et al., 2020), Spectronaut software v16.2 (Biognosys; Bruderer et al., 2015) and MaxQuant software v2.0.1.0 (MPI for Biochemistry; Sinitcyn et al., 2021). Protein identification was performed against the UniProtKB human and mouse reference proteomes (version 01/22), respectively, augmented by a set of 51 known laboratory contaminants, including the concatenated iRT peptide sequences. For all search engines, carbamidomethylation was set as a fixed modification, trypsin/P as the enzyme specificity, and a maximum of two missed cleavages were allowed. The minimum peptide length was set to seven amino acid residues, the maximum to 30. All FDR levels were set to a maximum of 0.01.

#### 2.6.1 DIA-NN Analysis

The search parameters of DIA-NN (version 1.8.1) were set to robust LC (high precision mode), library-free search enabled and library generation with smart profiling, deep learning-based spectra, neural network classified single-pass mode, cross normalization RT dependent with match between runs, use of isotopologues, no shared spectra, and heuristic protein and gene protein inference active. RTs and IMs prediction were enabled. When generating a spectral library, *in silico* predicted spectra were retained if deemed more reliable than experimental ones. Fixed-width center of each elution peak was used for quantification.

#### 2.6.2 MaxQuant TIMS-DIA Analysis

For data analysis in MaxQuant, the TIMS DIA algorithm v2.0.1.0 was used at default parameters. The MaxQuant discovery library *Homo sapiens* peptides, evidence, and msms files (release 2021-06-23) were used for searching. Protein quantification was based on both unique and razor peptides, FastLFQ was used for DIA quantification.

#### 2.6.3 Spectronaut Analysis

Search parameters in Spectronaut (version 16.2) were set as follows: analysis was performed in directDIA mode using default settings, except that the top 10 peptides per protein were used for quantification (major group top N).

#### 2.6.4 Generation of Annotated MS/MS Spectral Libraries

We built two annotated MS/MS spectral libraries based on experimental data and processing in DIA-NN software (version 1.8.1) using the search settings described above. For HEK293 cells, the spectral library was generated from the analysis of two non-fractionated lysates using both the short and long gradients described above with two technical injection replicates, plus single replicate files from the analysis of 35 BNE fractions generated for lysates P1 and P2, resulting in a total of 78 DIA-MS files. For the analysis of mouse cardiomyocytes, we chose a simplified library approach which included only non-fractionated measurements of one sample each from LVCM; RVCM and ACM. Each sample was analyzed using both the short and long gradients described above, and with two technical injection replicates.

#### 2.6.5 Statistical Analysis of Complexome Profiling Data Sets

mCP software was developed in R version 3.6 (R Core Team, 2022). All searches for the detection of PPCs were performed in dynamic mode by the mCP R package with default settings including a fixed-filter value equal to 0.81, 185 Monte Carlo simulations, and Pearson correlation methods (default).The only exception was the analysis presented in Supplementary Fig. 12, which was performed in *de novo* mode (dynamic = FALSE). PPCs were detected using a subset of the CORUM database (version 03/2022) filtered either for human (HEK293) or mouse (cardiomyocyte) entries, respectively. For the analysis of mouse cardiomyocytes, known mouse cardiac PPCs such as the calcium handling complexes (SERCA2A/RyR2/PLN), the SLMAP complex, and the IDH3G complex were added manually to generate a custom database, which is incorporated into the example data of the mCP R package. The molecular weight marker values used are indicated in supplementary material 2 (Table 1).

**Table 1.**
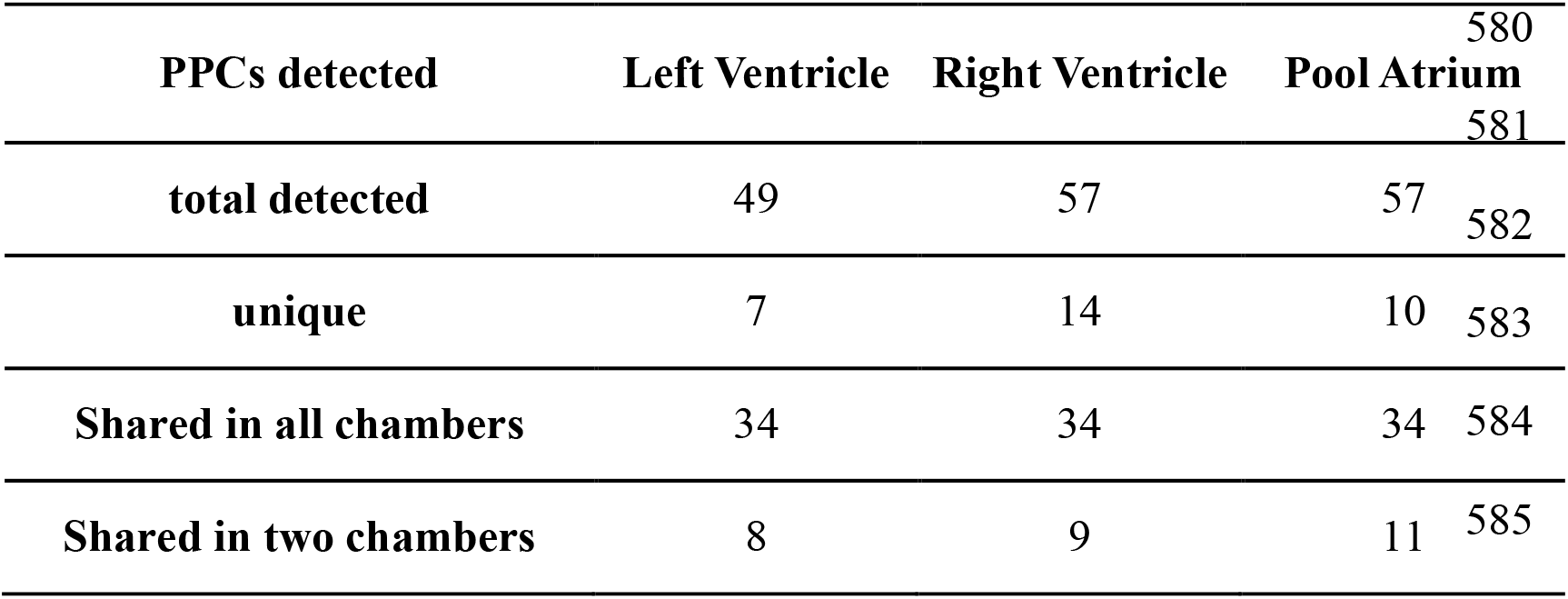
Protein/protein complexes (PPCs) detected in mouse heart chambers.

### 2.7 Additional Software Tools

Cartoon illustrations were created based on graphical components sourced from BioRender.com (publication agreement number PG26PW3SEG).

## 3 Results

### 3.1 Development and Benchmarking the mini-Complexome (mCP) Workflow

#### 3.1.1 mCP Workflow Design and Development

We explicitly designed the mCP workflow to be compatible with freely available and affordable consumables, starting with commercially available BNE mini-gels, which do not require significant optimization and afford excellent reproducibility (Fig. 1 A, Suppl. Fig. 8-10). Following non-denaturing electrophoresis, gel lanes are cut into 35 equidistant slices using an in-house manufactured stainless steel comb (Schmidt and Urlaub, 2009), alkylated and digested with trypsin. In-gel digestion of proteins is readily parallelized (e.g. in 96-well format (Schmidt et al., 2013)), and directly produces MS-compatible peptide mixtures. These samples are then analyzed using DIA-MS with short gradients (Bludau et al., 2023; Fossati et al., 2021a). The integration of CF-MS with DIA-MS has been demonstrated to produce highly reproducible and comprehensive complexome profiling data (Hay et al., 2023). The use of short gradients for DIA-MS acquisition leads to only a minor loss of information compared to longer gradient acquisitions, while also increasing throughput. We opted for a 20 min, or 72 samples per day, DIA-MS acquisition method, which allows for the analysis of two full BNE lanes per day and thus enables not only biological and technical replication but, more importantly, the comparison of different biological or disease states.

**Figure 1:**
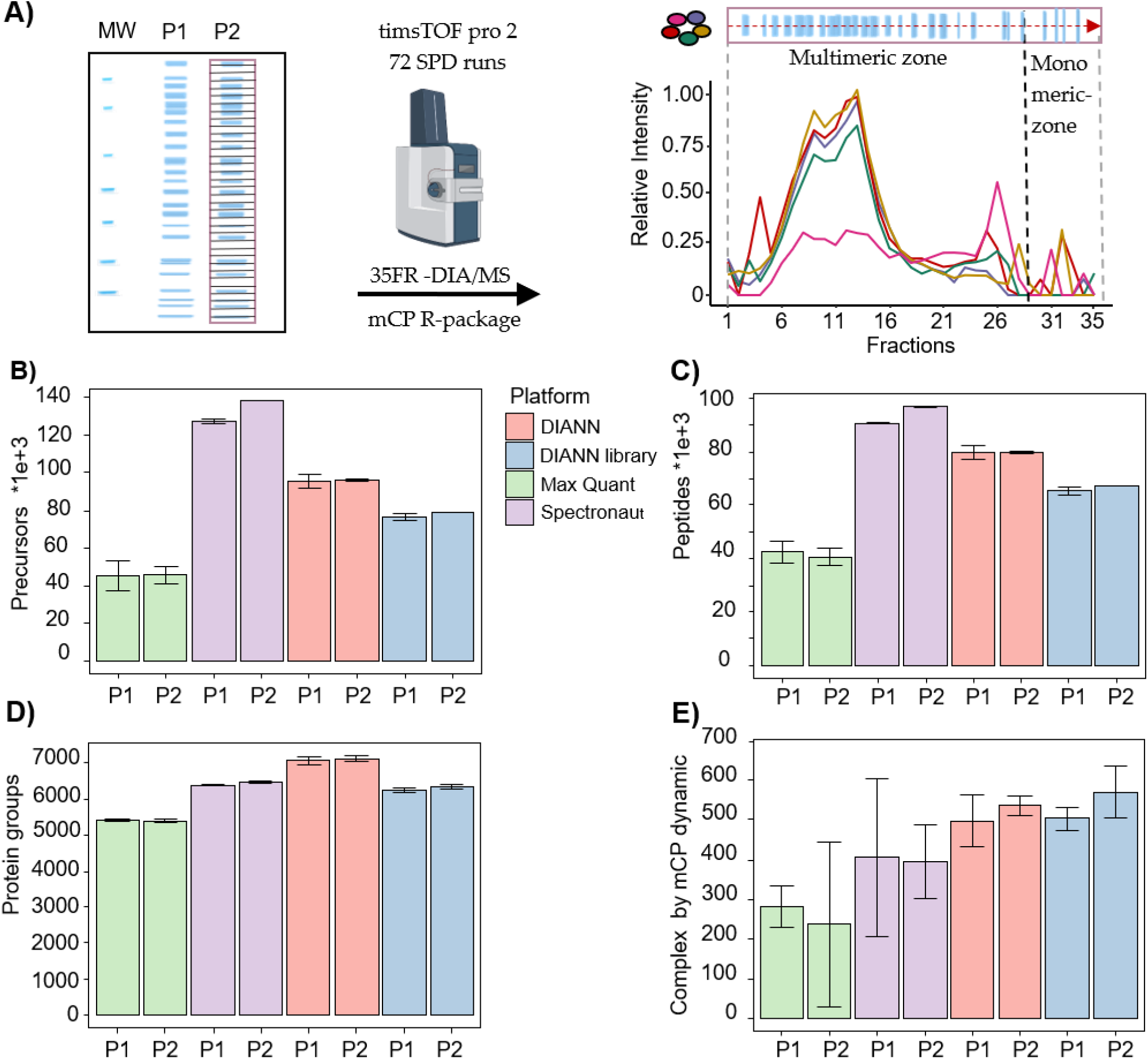
mCP workflow overview. A) Representative BNE-PAGE separation of HEK293 lysates. MW: Molecular weight marker. P1 and P2, biological replicates. Following fractionation to 35 slices per lane an in-gel digestion, slices were analyzed using 20 min DIA-MS acquisitions and their protein area values then plotted as a a function of fraction number. B), C), D) Performance comparison of different DIA-MS processing software packages on the detection of precursors, peptides, and protein groups from P1 and P2. E) Detection of protein complexes in these data sets using the mCP R package dynamic mode. Error bars represent 3 times the standard deviation of two technical replicates.

A second consideration for choosing a low-fractionation approach was to minimize protein input requirements. While high fraction workflows such as the one by Schulte et al. (Schulte et al., 2023) offer very high levels of detail, they also require significant amounts of starting material to yield at least low microgram protein amounts per fraction in order to support reproducible sample preparation and recovery. We aimed for a protein input amount ranging from 35-70 μg, which is consistent with the protein quantities recovered from human patient biopsies (Brandenburg et al., 2022) or single mouse organs. Assuming a roughly equal distribution of protein across the BNE, each of the 35 fractions generated in our approach will contain 1-2 μg of protein.

We tested the experimental workflow using 35 μg protein equivalents of HEK293 cell lysates. Following the workflow outlined above, we evaluated the performance of three widely used software packages for the analysis of the DIA-MS data, namely DIA-NN (Demichev et al., 2020), MaxQuant (Sinitcyn et al., 2021) and Spectronaut (Bruderer et al., 2015). The choice of processing software has been demonstrated to have a significant impact on the quality of DIA-MS experimental results (Fangfei Zhang et al., 2023; Fröhlich et al., 2022; Lou et al., 2023). We analyzed two biological replicates P1 and P2, each with two technical injection replicates (Supplementary 2 Fig. 8). All three software packages were employed for analysis without an a priori annotated MS/MS spectral library (so-called ‘directDIA’). In addition, we used a dedicated MS/MS spectral library for analysis by DIA-NN (‘DIANN library’). The results are summarized in Fig. 1 B-D.

In summary, we detected an average of 6.304 protein groups in the data set, evidenced by an average of 70.440 unique peptides and 87.999 precursors, respectively. On the precursor and peptide levels, Spectronaut significantly outperformed the other software packages, particularly MaxQuant (Fig. 1 B-C). Consolidation of the precursors and peptides into quantified protein groups showed more similar results, with DIA-NN in library-free mode showing the best results. This difference may hint at differences in the stringency of the protein grouping algorithms in the software.

#### 3.1.2 FDR-Controlled Processing of mCP Data Sets

We next set out to develop a bioinformatic tool for processing mCP data sets that enables automated detection of protein complexes under control of false discovery rates (FDR). Earlier CP-MS approaches mostly employed unsupervised approaches to data analysis such as hierarchical clustering (Giese et al., 2015), which, while powerful for comparative analysis, are subjective with regard to the parameters used, and lack FDR control. More recently, FDR-controlled CP-MS approaches have been developed that allow for an FDR control of PPC detections in CP-MS data sets by introducing an external ‘ground truth’, e.g. lists of PCCs obtained in comprehensive, well-curated PPC databases such as CORUM (Tsitsiridis et al., 2023) or String (Szklarczyk et al., 2023). Heusel et al. first published this approach and a corresponding computational tool named CCProfiler for the analysis of SEC-DIA-MS data sets (Heusel et al., 2019). Instead of the peptide-centric approach to data analysis inferred by building on large-scale peptide detection, this rather shifts the focus to a complex-centric approach where evidence for PPCs is extracted in a targeted manner, and FDR can be stringently assessed using target/decoy approaches.

Here we present a novel statistical approach to detect PPCs in CP-MS data sets at a controlled FDR, and its implementation in an R package called mCP which is optimized for low fractionation approaches (here, 35 fractions) as discussed above (Fig. 2). The FDR is assessed by a Monte Carlo simulation target decoy approach (MCs), where a series of decoys is generated and compared against PPC candidates in the experiment. We developed two modalities of search: a *de novo* search mode and a dynamic search mode. The *de novo* search calculates FDR under the assumption of independence in the Pearson’s coefficients (see Suppl. Material 1, Section 4, and Suppl. Material 3, Suppl. Fig. 12). The dynamic mode, on the other hand, prioritizes individual correlation within the PPC; it is more permissive and recommended for annotated protein complexes searches using CORUM or Complex Portal.

**Figure 2:**
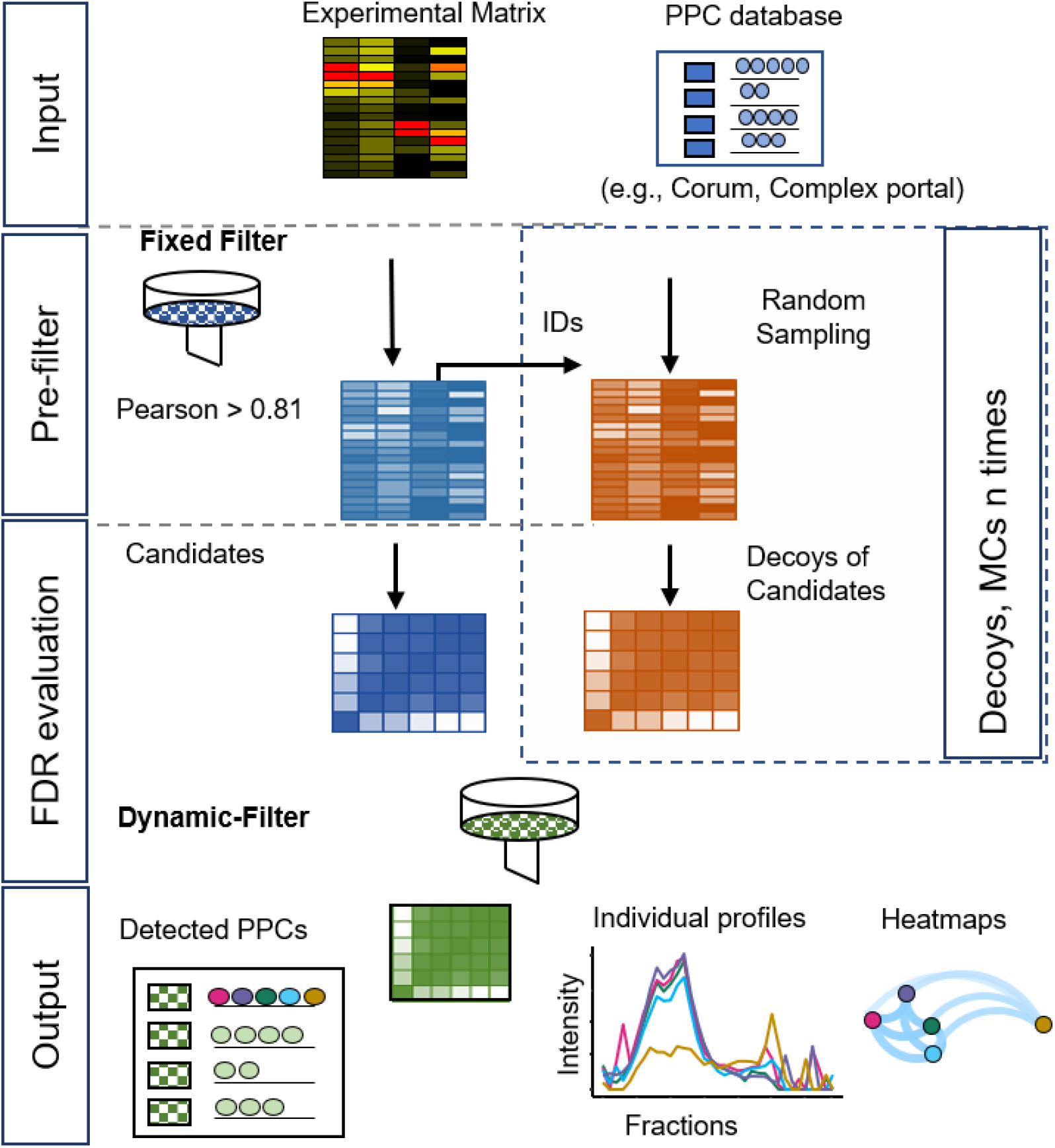
Schematic workflow for detection of PPCs using the mCP R package. The dynamic evaluation of protein complexes by the mCP-R package is divided into four stages: (i) the input of an experimental data matrix and a selected PPC database; (ii), pre-filtering of the data using empirically established thresholds and Monte Carlo simulations; (iii) estimation of the false discovery rate (FDR); and (iv) generation of graphical and table-based output formats.

In dynamic mode, mCP takes the experimental data matrix consisting of protein intensity values across the BNE-PAGE separation and a targeted list of protein complexes (from e.g. CORUM or ComplexPortal) and calculates Pearson’s correlation matrixes for each complex candidate present in both inputs. Then it applies a filter based on an empirically calibrated Pearson’s correlation value threshold. On the resulting candidates, it counts the number of hits and the average value of the Pearson correlation higher than the first filter, which is used as a dynamic Pearson filter specific for each protein complex to detect PPCs using a Monte Carlo simulation. In this step, random inputs from the experimental matrix are used as decoy matrices to establish false positives. Using our HEK293HEK293 test set described above, we found that the results of the Monte Carlo simulation stabilized after 185 iterations. Lastly, detected PPCs are reported to an FDR limit set by the user (see Supplementary Materials 1 and 3).

In the initial implementation of mCP, we observed over-assignment of protein complexes in the low apparent molecular weight (MW) regions of the BN-PAGE separated samples corresponding to apparent binary complexes consisting of low MW protein monomers. To address this, we implemented an additional ‘monomeric risk filter’ in the mCP function (see Suppl. Material 3).

Depending on the DIA processing algorithm used, our mCP approach detected an average of 426 (270-586) protein complexes in the HEK293 data set against the CORUM database, all at an FDR of 5% and with standard settings (Pearson’s filter 0.81, 185 Monte Carlo simulations) (see Suppl. Material 3). Among the different DIA data analysis softwares, DIA-NN with or without the use of a spectral library outperformed both Spectronaut and MaxQuant when it came to the detection of PPCs in this data set (Fig. 1D). While we cannot provide a detailed reason for this, we speculate that this may be a consequence of different stringencies in protein grouping, thus causing DIA-NN to present more protein complex constituent candidates. In spite of slightly lower performance compared to the library-free approach on the precursor, peptide, and protein group levels, DIA-NN processing using an experimental DIA library achieved the highest number of detected PPCs against the CORUM database (average 537 protein complexes, RSD 8.8%). We conclude that this approach is currently best suited for our workflow, which highlights the ongoing importance of experimental spectral libraries for select purposes of DIA-MS data analysis.

### 3.2 Benchmark Analysis of HEK293 Lysates

Using the optimized mCP workflow and parameters described above (Fig. 2; DIA-NN library input, Pearson’s filter 0.81, Monte Carlo 185 simulations), we next examined the results obtained on our HEK293 data set in more detail and benchmarked its performance against existing computational approaches like CCprofiler (Bludau et al., 2020; Heusel et al., 2019). HEK293 cells present an extremely well-characterized cell line model, with a significant number of PPCs contained in publicly available databases such as CORUM, STRING, or others.

mCP detected an average of 504±10 and 570±22 protein complexes in biological duplicates P1 and P2, respectively, demonstrating good technical reproducibility between replicate samples (Fig. 3A). We next applied the mCP algorithm to the HEK293 dataset published by Heusel et al. (Heusel et al., 2019) to test our computational approach on a higher degree fractionation (n=81), SEC-based data set of a comparable sample. Here, the mCP algorithm detected 490 protein complexes against the CORUM database, which stands up well to the 509 complexes detected by the authors (Fig. 3A). Moreover, mCP recovered 382/509 (75.0%) of the complexes detected with CCProfiler (Fig. 3B). Even among these two experimentally diverse datasets (our data, BN-PAGE, n=35 fractions; Heusel et al., SEC, n=81) we observed good overlap, with the two mCP replicates P1 and P2 describing 392/509 (77.0%) of the complexes detected by CCprofiler (Fig. 3B) (Bludau et al., 2020), underlining the robustness of our approach.

**Figure 3:**
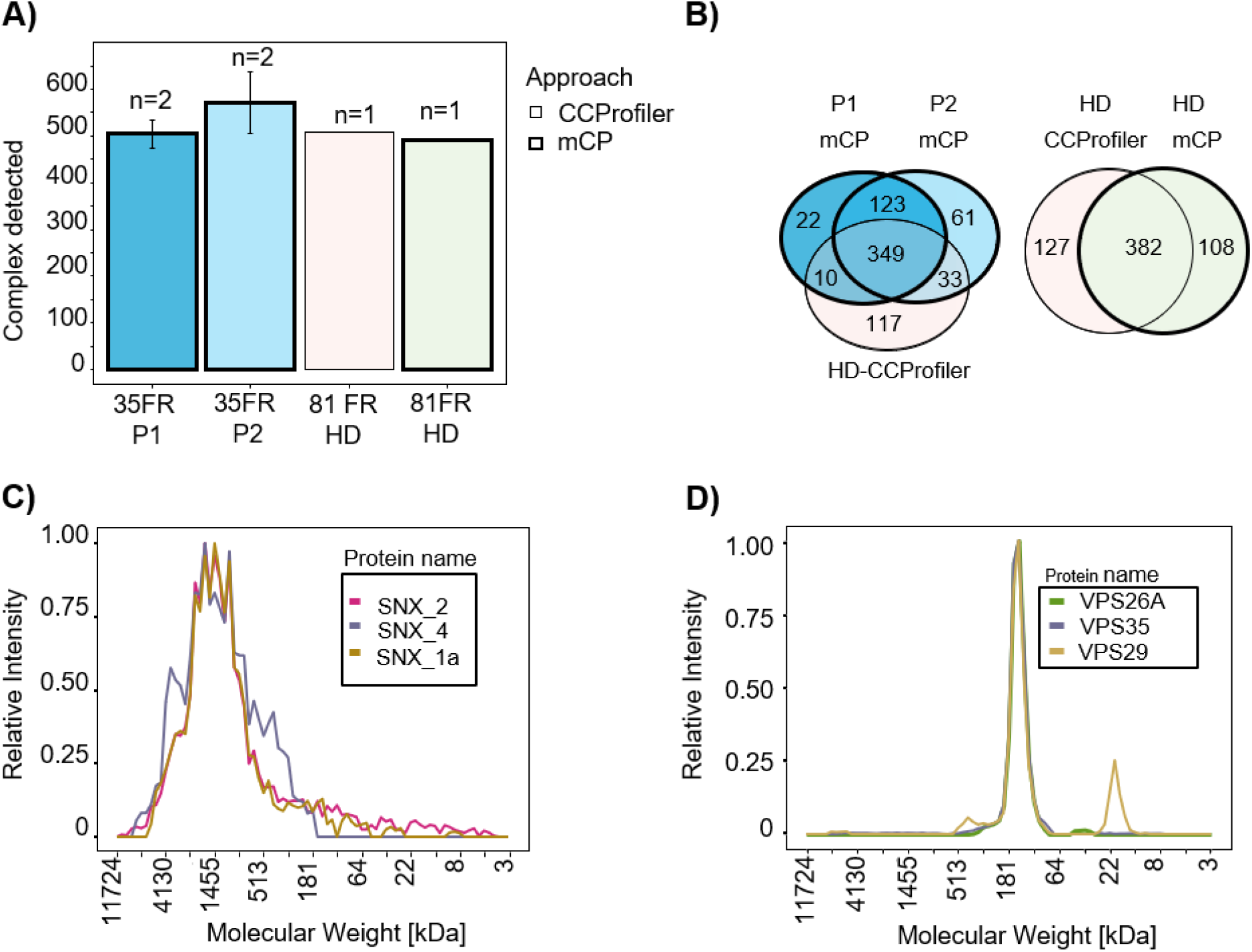
mCP workflow benchmarking for analysis of HEK293 cell lysates. A) Protein complexes detected in HEK293 cell lysates in our data set (P1 and P2, 35 fractions, 2 technical replicates) or in a reference dataset (81 fractions) (Heusel et al., 2019). Thick borders, analyzed with mCP; thin borders, analyzed with CCProfiler. B) Venn diagram of complexes detected by mCP R package using different data sets. mCP vs CCprofiler datasets analyzed by mCP (left panel) or CCprofiler dataset analyzed by CCprofiler and mCP (right panel). C), D) Elution profiles of SNX and VPS complexes exclusively detected by mCP in the 81 fraction reference data set.

Complexes detected exclusively by mCP in the 81 fraction reference data set include both complexes exhibiting wide, ‘hill-shaped’ elution profiles like a three-protein SNX complex, a membrane-associated complex involved in ribonucleoprotein assembly (Fig. 3C), as well as well-defined elution profiles like a VPS35-VPS29-VPS26A vacuolar sorting protein complex (Fig. 3D). While we cannot assess the completeness or purity of these complexes based on our data, our computational approach is clearly able to recover protein complexes not detected by other approaches from the same data.

### 3.3 Analysis of Individual Mouse Heart Cavities

Next, we set out to test our mCP workflow and computational approach on an altogether different sample type, i.e. the analysis of cardiomyocytes obtained from different cavities of an individual adult wild-type mouse heart in the C57BL6N genetic background. Previous work from both our lab (Foo et al., 2024) and other groups (Rosca et al., 2008; Gómez et al., 2009; Hou et al., 2019) has shown PCC and higher-order protein supercomplexes to play key roles in the heart, ranging from energy production (i.e. mitochondrial CI/III/IV supercomplexes) to Ca^+2^-handling (Alsina et al. 2019). While BN-PAGE has long been proposed as a tool to detect molecular defects in patients with disorders of oxidative phosphorylation (Van Coster et al., 2001), complexome profiling has so far been rarely applied either to the direct analysis of cardiac tissues, or to cardiomyocyte preparations of these. We thus investigated whether mCP can be applied to cardiac samples: specifically, cardiomyocytes isolated from different heart chambers (LV, RV, atria) from a single mouse. In many regards, these samples are more challenging than HEK293 cell lysates: (i) the limited protein amounts contained in cavities from individual animals preclude the use of native-like separations such as wide-bore SEC or wide-band BN-PAGE separations (e.g. as employed by Schulte et al. (Schulte et al., 2023)) combined with high fraction numbers, and (ii) publicly available PPC databases are largely derived from the analysis of human cancer research cell lines, while organ-specific proteomes from other species such as the mouse heart are strongly underrepresented e.g. CORUM 4.0 contains <1% of entries curated from tissue or cardiomyocyte samples (Foo et al., 2024).

A single heart was extracted from a 12 week-old wildtype (WT) male mouse and cardiomyocytes isolated by tissue lysis and sedimentation separately from the left and right ventricle (LV, RV) and the pooled atria together (PA). Proteins and protein complexes were fractionated by BNE and fractionated into 35 equidistant fractions (Supplementary Fig. 9 and 10). Again, all BNE slices were tryptically digested in-gel and analyzed by DIA-MS; the raw data were processed using DIA-NN with a library approach and evaluated using the mCP R package against the CORUM database as outlined above.

The results of protein complexes per chamber are shown in Table 1. Our exploratory analysis shows a total of 79 protein complexes found among all chambers, with 43% of the detected complexes (Table 1). The latter group comprises several well-described, high-abundance complexes such as the 20s proteasome, the immunoproteasome, the COP9 signalosome complex, and the CCT complex.

Interestingly, the mitochondrial protein complexes CI, CIII, and CIV, which are constituents of the CIx-CIIIx-CIVx respiratory chain supercomplex, showed markedly different migration profiles in the left and right ventricles (LV, RV) compared to the combined atria (Fig. 4). In particular, CI protein profiles presented a down-shift from an experimental apparent MW of 3.285 kDa in the LV and RV, which agrees with an expected MW of a CI2-CIII2-CIV2 supercomplex (Chavez et al., 2018) of ca. 1.165 kDa in mouse atria. Similarly, CIV proteins showed distributions to lower MW in the atrial samples, indicating a difference between respiratory chain supercomplex assembly states between the atria and ventricles in mouse, which to our best knowledge has not been documented so far (for analysis of whole mouse hearts see e.g. (Guo et al., 2017; Zheng et al., 2023)). Thus, we conclude that the mCP workflow here is sufficiently sensitive to perform analysis of cardiomyocytes from individual mouse heart cavities; and that it can visualize subtle compositional differences in protein complex assemblies through shifts in MW distributions.

**Figure 4:**
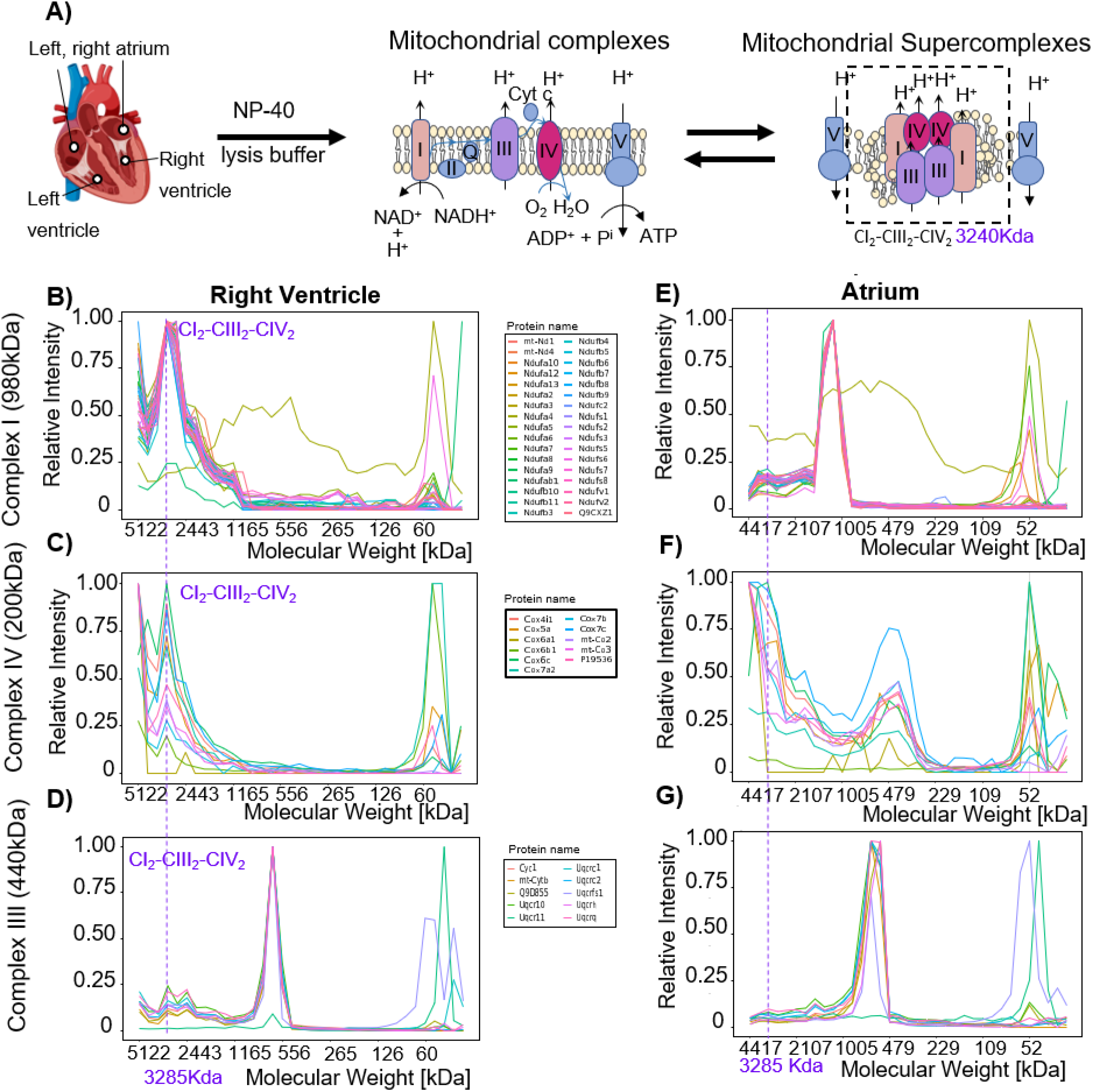
Detection of mitochondrial respiratory chain supercomplexes in cardiomyocytes. A. Schematic overview of mitochondrial respiratory chain (super-)complexes. B, E. Complex I profiles in the right ventricle (RV, left) and the atria (right). C, F. Complex IV profiles in the right ventricle (RV, left) and the atria (right). D, G. Complex III profiles in the right ventricle (RV, left) and the pool atria (right). The vertical dashed line in indicates the experimental apparent MW of the respiratory chain supercomplex at 3285 kDa.

## 4 Discussion

Here we present mCP: a complete experimental workflow for rapid, global complexome analysis coupled with an R package that allows for the detection and FDR control of PPCs. The mCP workflow consists of a BN-PAGE separation in a commercial minigel format; fractionation into 35 fractions and in-gel digestion; rapid protein abundance profiling across fractions by DIA-MS; protein identification by DIA-NN software; and finally, data analysis using the mCP R package.

mCP was designed to work on a reduced number of 35 fractions compared to most complexome profiling workflows. This idea was initially supported by the work of Skinnider and Foster 2021 (Skinnider and Foster, 2021), who performed a comprehensive reanalysis of 206 published complexome profiling datasets and demonstrated that the recovery of known protein complexes stabilized at a minimum of 35 fractions. Following this reasoning, reducing the number of fractions allows mCP to work with minimal requirements for the input material, as demonstrated by our analysis of cardiomyocyte preparations from individual mouse heart cavities, as well as minimizing the amount of mass spectrometry instrument time, which allows researchers to improve their experimental designs by way of replication or including additional biological controls and comparisons. Here, using a DIA-MS method with 20-minute turnaround time, or 72 samples per day, allowed us to analyze two full replicates of each BN-PAGE lane in a single day.

With regard to protein identification and quantification softwares to perform the initial protein identification and quantitation, we observed the best results when using DIA-NN software with a spectral library approach, which resulted in the highest number of protein complexes detected in an initial HEK293 cell lysate benchmarking experiment. However, our downstream data analysis pipeline also showed competitive results with e.g. Spectronaut, another widely available DIA-MS processing software, increasing researchers’ flexibility in adopting mCP for their needs.

The bioinformatic detection of protein complexes under false discovery control in these datasets is still challenging, not least because most target-decoy approaches to false discovery rate estimation are based on large search spaces and well-defined coelution profiles, which limits their effectiveness to data sets with high degrees of fractionation. Here, we propose a different approach by rather looking at whole chromatogram correlation matrices, calculating Pearson’s correlation, and then using Monte Carlo simulations to estimate false discovery (Fig. 2).

The mCP bioinformatics approach has two modalities. The dynamic search and the *de novo* search. The dynamic search is deeply studied in this publication; it performs similarly to CCprofiler, while the denovo search looks for integral detection of protein complexes with a deep search testing algorithm that is a formal FDR calculation. A comparison is provided in Suppl. Material 3, Suppl. Fig. 12. We see the application of this function in the context of *de novo* searches for the detection or validation of unknown PPC candidates, while the dynamic algorithm is strongly recommended for a search based on annotated protein complexes databases (CORUM or Complexportal). A great aspect of the mCP bioinformatics approach is the possibility of evaluating parallel datasets or parallel databases, as many as processing cores are available in the computer. This opens the possibility of validating inferred networks (Zahiri et al., 2020) or “blind testing” with a *de novo* search or parallel systems evaluation of known complexes (Fig. 1, Fig. 3, Table 1).

In HEK293 cell lysate benchmarking data, we achieved similar performance to a previously published complexome profiling workflow and software, SEC-SWATH and CCprofiler (Fig. 3) (Heusel et al., 2019). The major improvement observed with mCP analysis of both our data set and the dataset published by Heusel et al. is in the ability to detect protein complexes which do not show a highly defined, narrow MW range elution profile. As a whole chromatogram approach, mCP shows particularly improved performance for detecting broad, ‘hill-shaped’ elution profiles that are associated with protein complexes lacking a uniformly defined stoichiometry (Fig. 3C). We have begun to leverage this advantage for the analysis of very high MW, membrane-associated functional protein complexes that may exist as dynamic, non-covalent ‘functional rafts’ rather than as defined stoichiometric complexes (Foo et al., 2024). While it may be impossible to call out defined complex stoichiometries under these circumstances, differential analysis may nonetheless allow us to detect biochemical changes in large-scale complexome comparisons.

A major inherent limitation of mCP as a targeted data analysis approach lies in its dependence on the availability of comprehensive protein complex databases such as CORUM (Tsitsiridis et al., 2023) or Portal Complex (Meldal et al., 2015). The scope of the database effectively constrains the detection of proteins, as highlighted by the presence of 1753 human, but only 694 mouse protein complexes in CORUM as of March 2022. A large part of proteome research is directed towards cancer research in human disease models and backgrounds; the situation for other organ or disease backgrounds may be much less favorable. Going forward, this may be addressed by community efforts to increase protein complex coverage in these scenarios. As another technical limitation, the mCP R package currently contains March 2022 versions of CORUM and Portal Complex. Automated database updates in our R package for example, the CORUM database such as BioPlex R (Geistlinger et al., 2023) could not be made to work by us; therefore, an updated version of the database should be introduced by the user in our R package manually to do the search on the last database versions.

Apart from profile plots for each protein complex, mCP plots network heatmaps for each detected protein complex generated by the corrr package (Kuhn M et al., 2022). Network and heatmap plots are a good starting point to visualize the strength of interactions. We are currently evaluating the use of a more number-based approach to evaluate complex stability as well as compositional changes.

## 5 Conclusion

We developed a novel experimental and data processing workflow to streamline BN-PAGE-based complexome profiling analysis called miniComplexome Profiling, or mCP. mCP consists of a rapid experimental workflow based on readily available and affordable resources, as well as a novel bioinformatics tool that is able to detect known protein complexes at controlled false discovery rates. The reduced material requirements open the possibility of applying this workflow to sparse or rare samples, i.e. patient biopsies or organoids, extending the use of complexome profiling to perform differential analysis of e.g. multiple cellular states from a system biology perspective.

mCP is suitable for rapid screening of complexes to detect variations between different conditions. Using mCP, we detected different mitochondrial supercomplexes in RV vs. atrium. RV supercomplex stoichiometry correlates to a previous mouse supercomplex model (Chavez et al., 2018). Nevertheless, evaluation and comparison tools need to be developed to further detect alterations between datasets, there are already some other tools available (Meldal et al., 2015; Rizzetto et al., 2018). On a high number of fractions, we believe mCP could get further development to be able to identify unknown protein complexes.

## 6 Acknowledgements

HA would like to thank Phillip Schad for his contribution to the conceptualization and development of the mCP R package, as well as Resul Elgin (Georg August University Göttingen) and Jannis Anstatt (Max Planck Institute for Multidisciplinary Sciences) for testing software functionality. Mufassra Mushtaq, Brigitte Korff and Birgit Schumann (University Medical Center Göttingen, UMG) provided valuable help with the mouse experiments. CL would like to thank Jeanes Strebe (University Medical Center Göttingen) and Monika Raabe (Max Planck Institute for Multidisciplinary Sciences) for excellent technical assistance. Mass spectrometric analysis was supported by the UMG Core Facility Proteomics.

## 7 Author Contributions

Conceptualization, HA and CL; methodology, H.A, LN and CL; animal documentation and sample harvesting, BF; investigation, HA and BF; formal analysis, HA; software and scripting, HA and NP; writing and editing, HA and CL; revision of the document, CL, HU, and SEL. All authors have read and agreed to the published version of the manuscript.

## 10 Data availability

The mass spectrometry proteomics data have been deposited to the ProteomeXchange Consortium via the PRIDE (Perez-Riverol et al., 2021) partner repository with the dataset identifiers PXD049340 and PXD049426. The mCP R package is available at https://github.com/hugoagno3/mCP.

