## Supplementary Materials 1, 2, and 3. for "mini-Complexome Profiling (mCP), an FDR-controlled workflow for global targeted detection of protein complexes"

### *Supplementary Material 1: mCP Script*

This document provides a short tutorial to perform mCP analysis using the mCP R package. The tutorial is also available on the following link: https://github.com/hugoagno3/mCP.

#### 1 Overview

The mCP program provides a set of functions for targeted protein complex detection from co-fractionation mass spectrometry-based proteomics experiments. It is designed to work with experimental data in the form of protein abundance matrices. The program identifies protein complexes present in the experimental data using an external protein complexes database (e.g. CORUM or Complex Portal) as a reference. It then plots profile (Suppl. Fig. 1), heatmap (Suppl. Fig. 2), and network plots (Suppl. Fig. 3) for each detected protein complex. The R package contains experimental co-fractionation data from human HEK293 cell lysates and from mouse heart cardiomyocytes as examples.

This vignette provides a step-by-step guide to using mCP to analyze proteomics data and identify protein complexes of interest. In this tutorial, we will analyze a single co-fractionation experiment derived from HEK293 cell lysates fractionated by BNE-PAGE into 35 fractions. In addition, we provide a detailed explanation of the R package functions at the end of this tutorial.

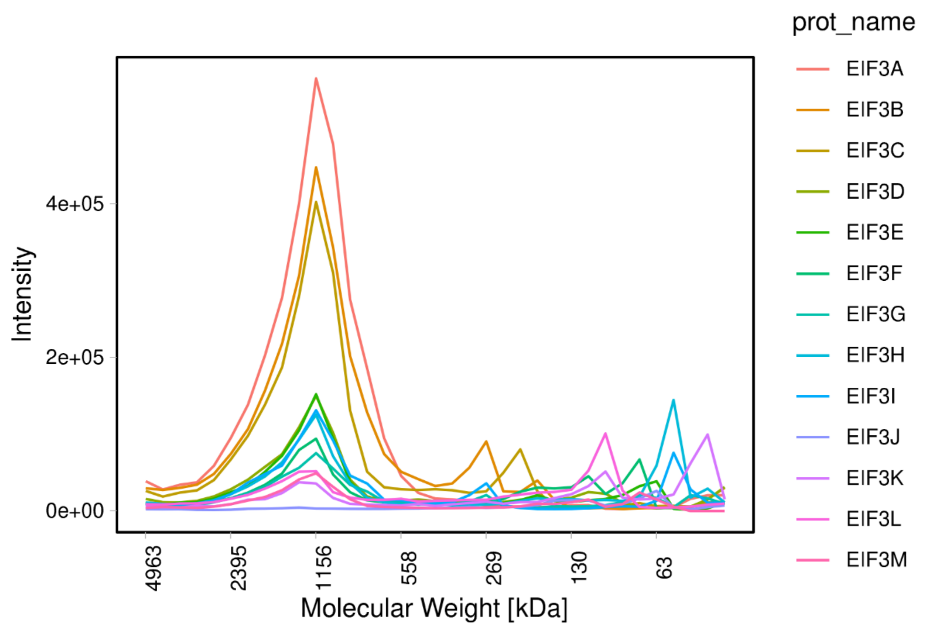


**Supplementary Figure 1: mCP R package example of complex profile detection of eIF3 complex.** Complex intensity profile as a function of molecular weight calibrated fractions.

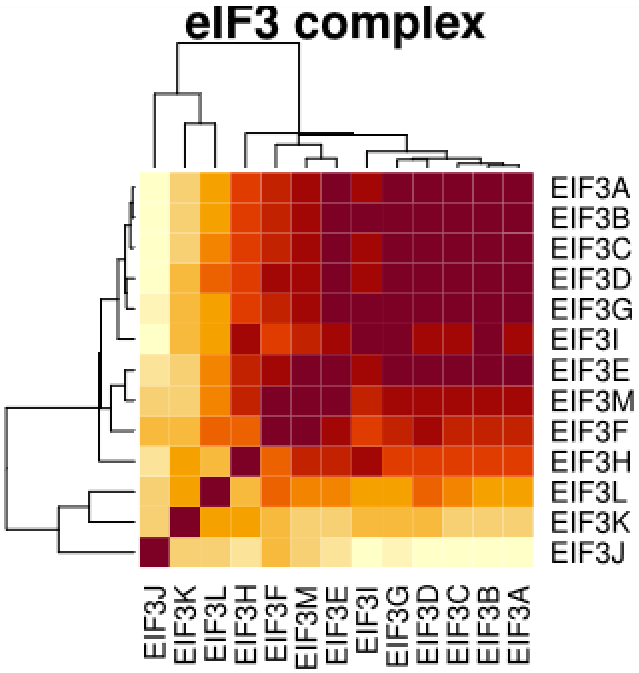


**Supplementary Figure 2: mCP R package example of Heatmap of Complex eIF3.** Pearson correlation matrix of protein components of eIF3 complexes, detected in 35 fractions. Range of colors white: 0 to dark brown: 1.

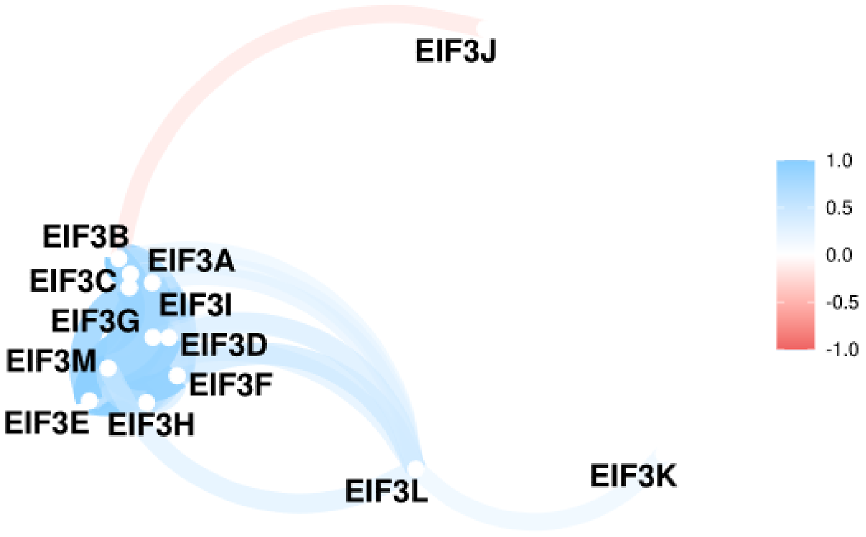


**Supplementary Figure 3: mCP R package example of Heatmap network of Complex eIF3.** A network representation from the Pearson correlation matrix of all protein components of eIF3 complexes, detected in 35 fractions. Range of colors white: 0 to dark brown: 1.

#### 2 Installation

To install the mCP package from GitHub, run the following code:

- ➢ library (devtools)
- ➢ devtools:install_github(“hugoagno3/mCP”)

#### 3 Processing instructions

##### 3.1 Data input

In this tutorial, we will use the Corum_Humans_Database file as an external protein complex reference database, and the Hek293_P2_1 file as experimental input. To perform targeted protein complex detection, only these two input files are required:

• A CORUM database of protein complexes (or a targeted list of interest) data frame with 3 columns: col1= "complex_id", col2= "complex_name" and col3= "protein_id".

• An experiment file data.frame with your experimental results in wide format. The first column is called "protein_id", with a single UniProtKB accession per row as elements. The rest of the columns contain protein intensities detected in each fraction of the co-fractionation experiment. The column's names of the fractions. It can be numeric names from 1 to the last number of fractions. For example, 1,2,3, […] 35 for 35 fractions, meaning 35 columns.

### Open the Corum protein complex database file

- ➢ data(Corum_Humans_Database)

### Open the experimental data files

- ➢ data(Hek293_P2_1)

Note: The mCP R package is focused on the detection of protein complexes, and it accepts only protein data level as an input matrix. The mass spectrometry data acquisition can be performed using data-dependent acquisition (DDA) or data-independent acquisition (DIA) modes.

#### 3.2 Data processing

To process the input data, there are two options. Option 1 – running the mCP function, or option 2 – running all three functions provided by the mCP package.

##### 3.2.1 Option 1: Running the MCP function

mCP() is an integrated function of mCP R-package. It requires an experimental protein abundance matrix dataset as input, and returns the following:

• A curated list of identified protein complexes, accompanied by a controlled False Discovery Rate filter (FDR). Each detected protein complex is an output element organized into the resulting list output of mCP.

The output list is structured with four distinct elements:

1. One fraction profile plot (absolute or relative abundance of each protein vs. fraction).

2. The number of intra binary interactions (protein-protein) with a Pearson correlation value higher than the filter (number of total binary hits) within pair components of each protein complex in the experiment.

3. The id of proteins of binary hits.

4. A heatmaps_seaborn of known protein complexes detected in the protein complexes database. In this tutorial, we use the CORUM database (http://mips.helmholtz-muenchen.de/corum/).

The above data is output in the following format:

1. pdf file with fraction-profile plots (absolute or relative abundance of each protein vs. fraction) of each potential protein complex to be detected. Before the FDR analysis.

2. pdf with heatmaps of potential candidates from 1 (before the FDR analysis).

3. pdf file with the detected protein complex profiles with a controlled FDR (this is analogous to 1, but without protein complexes, excluded based on FDR evaluation).

4. pdf with heatmaps of the detected protein complexes with a controlled FDR (this is analogous to 2, but without protein complexes excluded based on FDR evaluation).

5. A text file with log run information, and runs parameters.

6. CVS file containing all protein complexes detected, hits of binary interactions inside the protein complexes, and FDR detected by MonteCarloSimulation. An example can be found here:

```
# Read the Corum protein complex database file
➢ library(mCP)
➢ data(Corum_Humans_Database)
```

### Read the experiment files

- data(Hek293_P2_1)

###### Example #####

- mCP_Hek_P2_1 <- mCP(corum_database = Corum_Humans_Database,

```
experiment_data = Hek293_P2_1,
N_fractions = 35,
specie = "hsapiens",
method_cor = "pearson",
heatmap_seaborn = TRUE,
format = "pdf",
output_name = "m_CP_analysis",
filter = 0.81,
heat_map = TRUE,
relative = FALSE,
fdr_limit = 0.05,
n_simulations= 9,
monomeric_filter = FALSE,
Risk_fraction = 31,
dynamic = TRUE,
set_seed = TRUE,
display_weights = TRUE,
standard_weights = list(list(x =6, label= "2700 KDa"),
                        list(x = 11, label ="950 KDa"),
                        list(x = 14, label = "750 KDa"),
                        list(x =27, label ="146 KDa"),
                        list(x =30, label ="60 KDa")))
```

##### 3.2.1.1 Outputs plots per Protein complex detected

- mCP_TEND_out_Hek_P2_1_teste_11$Respiratory chain complex I (intermediate VII/650kD), mitochondrial

Output 1: [[1]]

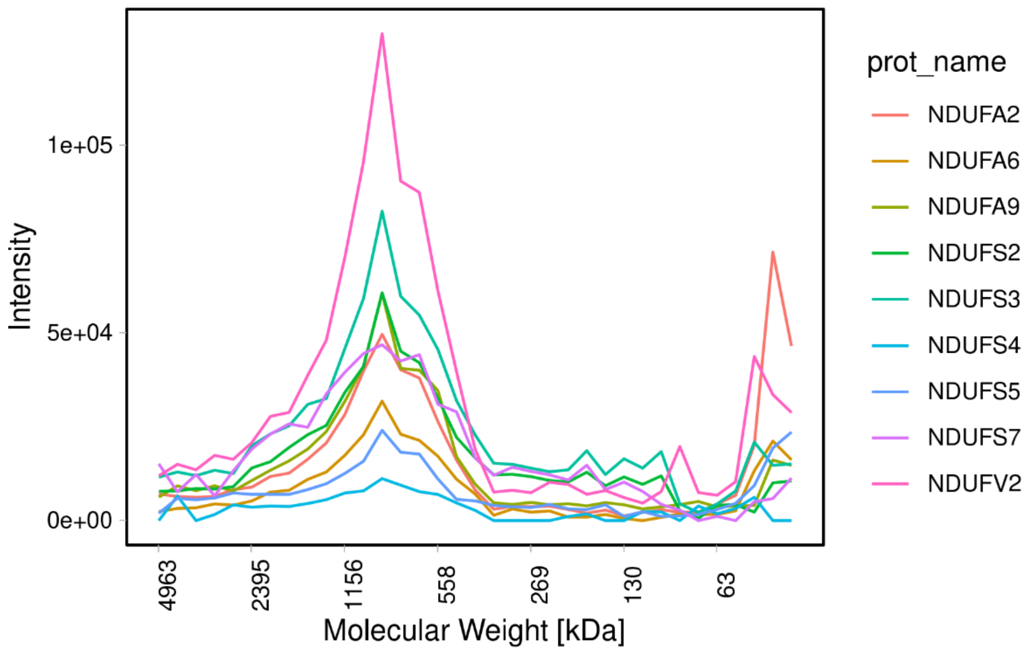


**Supplementary Figure 4: Output Complex profile plot of Respiratory chain complex I (intermediate VII/650kD).** Complex profile Intensity detected in the MS vs. molecular weight detections.

Output 2-4:

[[2]] Hits 1 9

[[3]] [[3]]$interactions [1] "NDUFS5:NDUFA6" "NDUFA6:NDUFA9" "NDUFS3:NDUFA9" "NDUFA9:NDUFS2" "NDUFS7:NDUFS2" "NDUFS2:NDUFS3" [7] "NDUFA6:NDUFS5" "NDUFA6:NDUFV2" "NDUFS3:NDUFV2"

[[3]]$Pearson [1] 0.9434851 0.9346499 0.9487094 0.9306028 0.9495674 0.9727912 0.9434851 0.9427735 0.9544865

[[4]]

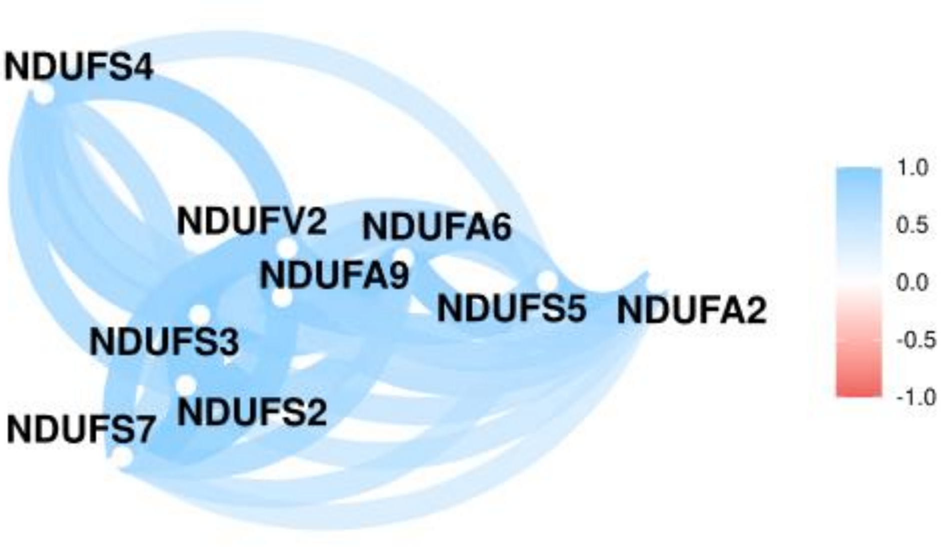


**Supplementary Figure 5: Output network heatmap plot of respiratory chain complex I (intermediate VII/650kD).** A network representation of Pearson correlation matrix of all proteins in the respiratory chain complex I (intermediate VII/650kD).

Note: mCP has two possible evaluation algorithms to detect false positive that can be activated by the parameter dynamic = TRUE or FALSE. Explained in section 4.

##### 3.2.2 Option 2: Run mCP function individually

The following code guides you to get the output step-by-step:

- library(dplyr)
- library(mCP)

```
# Read the Corum protein complex database file
```

- data(Corum_Humans_Database)

```
# Read the experiment files
```

- data(Hek293_P2_1)

```
#### RUN mCP list that creates a list of potential protein complexes
```

```
CL_hek_P2_1<- mcp_list(corum_database =  Corum_Humans_Database,
        experiment_data = Hek293_P2_1,
        N_fractions = 35,
        specie = "hsapiens",
```

method_cor = "pearson",

heatmap_seaborn = TRUE)

###### Run the output of mcp_list into the cpp_ploter function.

- out_Hek_P2_1 <- cpp_plotter(complex_list = CL_hek_P2_1,

format = "pdf",

output_name = "m_CP_analysis",

filter = 0.81,

N_fractions = 35,

heat_map = TRUE,

relative = FALSE,

display_weights = TRUE,

standard_weights = list(list(x =11, label= "1049KDa"),

list(x = 13, label ="720 KDa")))

###### Run the Experimental matrix and the output of cpp_ploter into the fdr_mcp function. Keep the same filter value!

- FDR_DIANN_dDIA_Hek_P2_1_<- fdr_mCP(corum_database= Corum_Humans_Database,

Output_cpp_plotter = out_Hek_P2_1,

experiment_data=Hek293_P2_1,

file_name = "m_CP_analysis",

N_fractions = 35,

specie = "hsapiens",

fdr_limit = 0.05,

filter=0.81,

Risk_fraction = 31,

monomeric_filter = FALSE,

set_seed = TRUE,

n_simulations= 185)

Note: This function will perform an FDR filter and simulation. It is important to do the simulation with the same filter as the previous function. If the filter value varies, the simulation loses its meaningfulness. If you want to run the *de novo* search, use the same code but the function fdr_standard_modified() instead of fdr_mCP ().

The last step to get the protein complexes filtered by FDR is to get back to the list and run cpp_plotter again.

Note: If you wish to have the plots of these last FDR protein complexes, you could filter the names of the protein complexes detected into the first list (from mcp_list) and run cpp_ploter again.

```
#### list to plot
```

- CL_final_output<- CL_hek_P2_1[names(FDR_DIANN_dDIA_Hek_P2_1_)]

```
#### Then run again this list on the cpp_plotter function as follows.
```

- out_Hek_P2_1_final_output <- cpp_plotter(complex_list = CL_final_output,

```
                format = "pdf",
                output_name = "m_CP_analysis",
                filter = 0.81,
                N_fractions = 35,
                heat_map = TRUE,
                relative = FALSE,
                display_weights = TRUE,
                standard_weights = list(list(x =6, label= "2700 KDa"),
                                        list(x = 11, label ="950 KDa"),
                                        list(x = 14, label = "750 KDa"),
                                        list(x =27, label ="146 KDa"),
                                        list(x =30, label ="60 KDa")))
```

###### 3.2.2.1 Output examples.

The output list out_Hek_P2_1_final_output has a list of protein complexes. Each protein complex is an element of that list with 4 objects.

1. The first object is a profile of proteins, fractions vs. intensities.

➢ out_Hek_P2_1_final_output[["13S condensin complex;Condensin I complex"]][[1]]

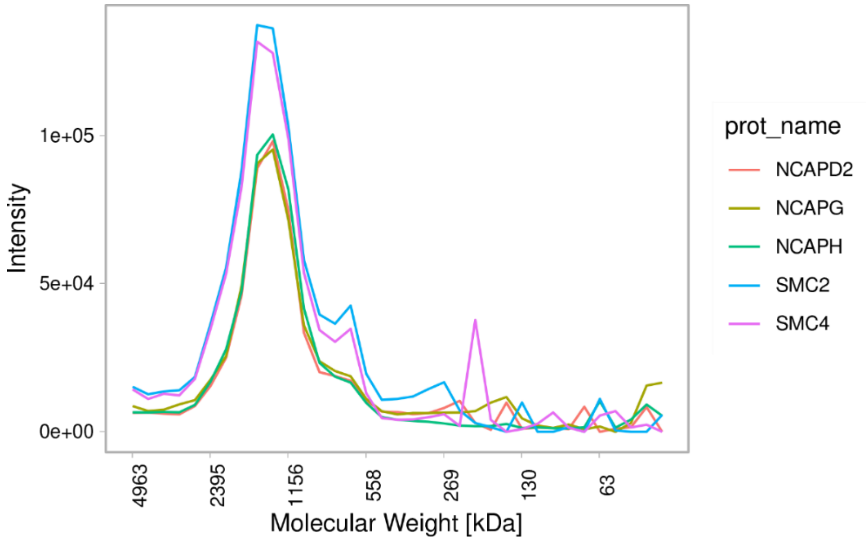


**Supplementary Figure 6: Output element 1, complex profile of complex profile detection 13S condensin complex; Condensin I complex.** Complex intensity profile detected in the MS vs. molecular weight calibration.

2. The second object is the number of hits detected in the experiment (the count of the significant binary interactions). So, the binary Pearson's correlation values in the experiment that were greater than a specified threshold.

➢ out_Hek_P2_1_final_output[["13S condensin complex;Condensin I complex"]][[2]]

```
[[2]]
  Hits
1   10
```

3. The name of the proteins involved in the binary interaction detected (experimental protein-binary Pearson correlation higher than the filter) and its Pearson’s correlation of the significant binary interactions.

➢ out_Hek_P2_1_final_output[["13S condensin complex;Condensin I complex"]][[3]]

[[3]]

[[3]]$interactions

[1] "NCAPH:NCAPD2" "SMC4:NCAPD2" "NCAPH:NCAPG" "SMC4:NCAPG" "NCAPG:NCAPH" "SMC4:NCAPH"

[7] "NCAPG:SMC2" "SMC4:SMC2" "NCAPG:SMC4" "SMC2:SMC4"

[[3]]$Pearson

[1] 0.9939944 0.9552057 0.9912144 0.9463247 0.9912144 0.9585936 0.9776026 0.9662482 0.9463247 0.9662482

4. Network heatmap constructed from the correlation matrix.

➢ out_Hek_P2_1_final_output[["13S condensin complex;Condensin I complex"]][[4]]

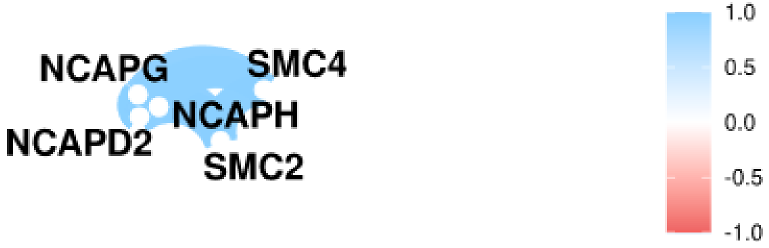


**Supplementary Figure 7: Output network heatmap 13S condensin complex; Condensin I complex.** A network representation of a Pearson's correlation matrix of all proteins in the 13S condensin complex; Condensin I complex.

##### 3.3 Data Processing (Input of matrix with protein_id as first column and fractions)

In a typical co-fractionation experiment, samples are measured in duplicate. The mCP R package provides a function to deal with the increased number of columns. The function input can be a data.frame imported by the following function read.table( ).

➢ NAmatrix_P2_1 <- read.table("Hek293_P2_1.csv",sep =",",dec = ".", header= T)

Once imported, you will get this file, we provide it in the R package, just write:

➢ data(NAmatrix_P2_1)

The function calc_mean_matrix will recognize the replicates by a tag like *A* and *B* as part of the names for the MS file and the *Frac_index*. It is the position in which you can find the number of fractions in the column name, separated by underscores. For example, the frac_index=5 is date_Surname_Measurement_A_01 because the number for the fraction is 5 spaces between underscores, like a_b_c_d_FractionNumber.

##### Note: Here the frac_index= is 17. and the pattern_group= _A_ and pattern_group= _B_

- names(Hek293_P2_1[,2])

[1] "D:\\Current rojects\\Projects_2022\\2022_77_H_Amede_mCP_Hek293_Cells\\03_Raw Data timsTOF Pro\\P2\\H_Amedei_04112022_P31_HEK293_P2_1_A_01_GA1_1_6588.d"

names(Hek293_P2_1[,37])

[1] "D:\\Current Projects\\Projects_2022\\2022_77_H_Amede_mCP_Hek293_Cells\\03_Raw Data timsTOF Pro\\P2\\H_Amedei_04112022_P31_HEK293_P2_1_B_01_GA1_1_6626.d"

So we can run the function as follows.

```
Hek_293_MA_P2_1 <- Calc_mean_matrix(NAmatrix = NAmatrix_P2_1,
                   pattern_group_A = "_A_",
                   pattern_group_B = "_B_",
                   frac_index=17,
                   Protein_ID_column = 1,
                   save_file = TRUE,
                   save_name = "Hek293_P2_1.csv")
```

Note: The user can also use their own script to generate the average of replicate experiments. This function is currently limited to accommodating two replicates. The mCP function does not accept NA values. All NA values in our case are replaced by 0.

#### 4 mCP main functions

The mCP package provides the following main functions:

**mCP():** This function is an integrated function of the mCP R package that needs as input experimental data plus a target list, and returns a list of plots, binary total hits, ids of proteins of binary hits, and heatmaps_seaborn of known protein complexes detected in the target list (for example, in the CORUM database). It has two search modalities: dynamic (dynamic = TRUE) and *de novo* (dynamic = FALSE). The parameter dynamic = TRUE activates the dynamic search function by using in the simulation the function fdr_mCP, while the parameter dynamic = FALSE, will use the fdr_mCP_standard_modified () function. The dynamic search uses a Pearson's correlation coefficient filter prior to the generation of decoys. It is optimized by the average of Pearson's coefficient higher than a threshold detected by the filter. While the *de novo* search assures the decoy generation in every simulation by performing two parallel simulations and calculating the false discovery rate under the assumption of independence of Pearson's coefficient. We recommend the dynamic = FALSE for *de novo* detection of protein complexes since it is a robust formal detection of FDR. For exploratory purposes, with a list of known and validated protein complexes like (the CORUM or Complex Portal databases), the dynamic = TRUE is recommended.

**mcp_fdr():** Is the function employed in the dynamic search. This function performs a serial evaluation of false positives in a single simulation. It first compares connections within PPCs; connections are essentially binary interactions (yes/no interactions) that have a Pearson's correlation coefficient higher than a certain threshold. A correlation matrix is calculated per PPC, and a filter value is set. The number of connections is compared to the decoys connections and if the decoys connections are the same or higher, it is considered a false positive. The filter value to detect a connection is the average Pearson's value detected per PPC on the target matrix, which is higher than the filter value.

**mcp_fdr_standard_modified():** Is the search performed on mCP function when dynamic = FALSE. This function calculates the FDR under the assumption of independence in the Pearson's correlation coefficient. It performs two independent simulation rounds. The first simulation compares connections within PPCs; connections are essentially binary interactions (yes/no interactions) that have a Pearson's correlation coefficient higher than a certain threshold. In simulation 1, a correlation matrix is calculated per PPC, and a filter value is set. The number of connections is compared to the decoy's connections and if the decoys connections are the same or higher, it is considered a false positive. The filter value to detect a connection is the minimum Pearson's value detected per PPC higher than the filter value. The second simulation compares the average of the same top connections of Pearson's correlations between target and decoys. The number of connections is defined by the connections detected by the filter value in the function. A false positive is assigned to a decoy if the Pearson's top connections average is equal to or higher than the target. For each PPC, the FDR is calculated as the sum of total false positive divided by the sum of simulations (simulation 1 + simulation 2).

**Calc_mean_matrix():** pre-processes the data before running mcp_list().

**mcp_list():** Extracts potential protein complexes from the experimental dataset, using the protein complexes database as a reference. This function extracts the intensities of all protein complexes from the protein complexes database (CORUM or complex portal) and generates a list of all potential protein complexes present in the experimental dataset. Each element of the list is identified as a potential protein complex and named with its corresponding complex name. Each protein complex element contains two elements:

*my_protein_complex_mcp_list[1][1]* is a dataframe composed of the Id proteins of the proteins, names, and fractions and their corresponding intensities per fraction. Names are annotated to the UniProtKB accession ids by a repository R package called gprofiler2.

*my_protein_complex_mcp_list[1][2]* is a correlation matrix of these elements, this correlation matrix is done by Pearson's correlations (but Kendall and Spearman algorithms are also possible in that step). The matrix was calculated as a correlation matrix of all detected components of each detected protein complex.

The list of potential protein complexes is the input for the next function called **cpp_plotter().** This is the pre-search stage. Where protein complexes composed of at least two components with intensities different from cero are selected. Then the first filter is run: The filter is based on a minimum definition of protein complex. So in this step, it keeps only protein complexes with at least one significant coelution hit (a binary interaction with a Pearson's correlation higher than the filter).

**cpp_plotter():** The cpp_plotter() function takes a list of potential protein complexes as input. The function first filters out any complexes composed of less than two components with non-zero intensities. Then, it applies a filter based on the minimum definition of a protein complex, keeping only complexes with at least one significant co-elution hit. The function also filters the list of complexes based on the presence of co-eluting proteins, requiring at least one binary interaction higher than the filter for proteins within the candidate complex. Finally, cpp_plotter() generates complexome profiling plots, heatmaps, and network plots of the proteins within the selected complexes.

**cpp_plt_sim():** Is an intermediate function used in Monte Carlo simulations fdr_mCP(). The function is the same as cpp_plotter(), but it is prepared to use a vector of different filters according to the average of the Pearson's correlation calculated for each protein complex (higher than the first filter). The function requires a list of potential protein complexes as input. The function first filters out any complexes composed of less than two components with non-zero intensities. Then, it applies a filter based on the minimum definition of a protein complex, keeping only complexes with at least one significant co-elution hit. The function also filters the list of complexes based on the presence of co-eluting proteins, requiring at least one binary interaction higher than the filter for proteins within the candidate complex. Finally, cpp_plt_sim() generates complexome profiling plots, heatmaps, and network plots of the proteins within the selected complexes.

### *Supplementary Material 2: Molecular Weight Calibration*

This document describes the molecular weight calibration performed for the BNE-PAGE separations shown in the main manuscript. The calibration is based on the unstained Native molecular weight ladder (NativeMark, Invitrogen, catalog number LC0725; Supplementary Table 1).

| Marker MW (kDa) | Fraction # Suppl. Fig. 8 | Fraction # Suppl. Fig. 9 | Fraction # Suppl. Fig. 10 |
|---|---|---|---|
| 1048 | 12 | 11 | 10 |
| 720 | 15 | 14 | 13 |
| 480 | 19 | 18 | 17 |
| 242 | 23 | 22 | 21 |
| 146 | 27 | 26 | 25 |
| 60 | 30 | 29 | 28 |

**Supplementary Table 1: Molecular weight marker information for BNE-PAGE calibration**

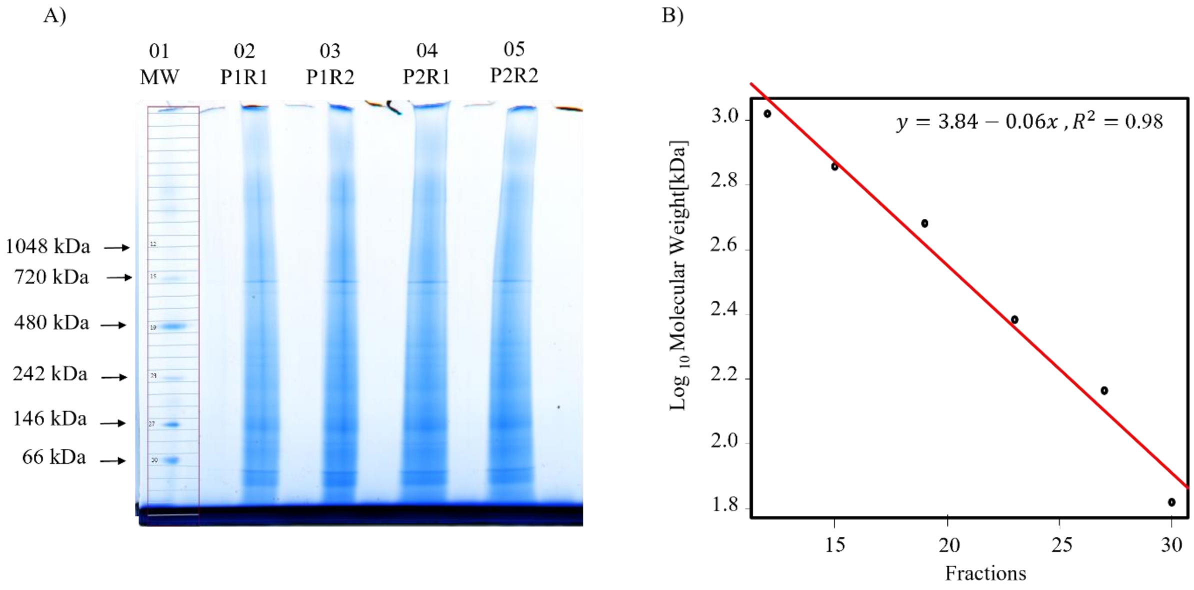


**Supplementary Figure 8: Molecular weight calibration of HEK293 lysate using BNE-PAGE** A) Scan of BNE-PAGE 3-12% native tris gel stained by colloidal Coomassie Blue. Lane 01, molecular weight marker (MW). Lanes 02 and 03: technical replicates of HEK293 biological replicate 1. Lanes 04 and 05, technical replicates of HEK293 cells biological replicate 2. B) $Log_{10}$ linear regression of lane 1 marker molecular weight (MW) vs observed fraction numbers.

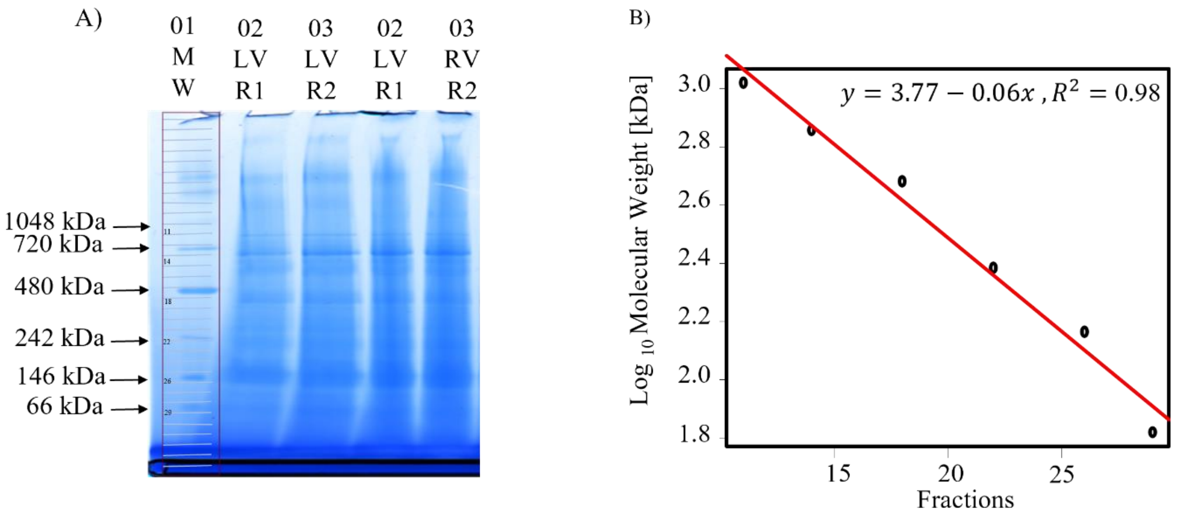


**Supplementary Figure 9: Molecular weight calibration of cardiomyocyte lysate from left and right mouse heart ventricle using BNE-PAGE.** A) Scan of BNE-PAGE 3-12% native tris gel stained by colloidal Coomassie Blue. Lane 01, molecular weight marker (MW). Lanes 02 and 03, technical replicates of left ventricle cardiomyocytes. Lanes 04 and 05, technical replicates of right ventricle cardiomyocytes B) $Log_{10}$ linear regression of lane 1 marker MW vs observed fractions numbers.

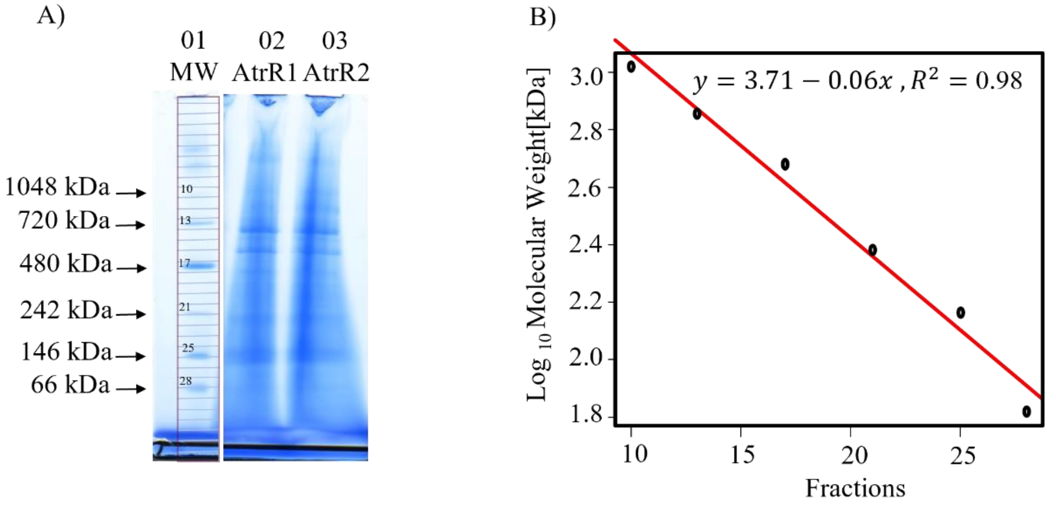


**Supplementary Figure 10: Molecular weight calibration of cardiomyocyte lysate from mouse heart atrium using BNE-PAGE.** A) Scan of BNE-PAGE 3-12% native tris gel, stained by colloidal Coomassie Blue. Lane 01, molecular weight marker (MW). Lanes 02 and 03: technical replicates of atrial cardiomyocytes. B) $Log_{10}$ linear regression of lane 1 amerrker MW vs observed fraction numbers.

### *Supplementary Material 3: mCP Bioinformatic Approach*

This document specific information about the mCP R package's bioinformatic approach for the global detection of annotated protein complexes from Complexome Profiling (CP) data.

#### 1 The mCP R package

The bioinformatic approach is specifically tuned for CP experiments with a lower number of fractions (n<40) while maintaiing FDR control using a Monte-Carlo simulation approach.

##### 1.1 mCP - Inputs

Firstly, mCP requires only two inputs to perform protein complexes detection, the experimental matrix in wide format and the target list of protein complexes. It then creates a list of candidate protein complexes according to a target list (CORUM or Complex Portal). Next, it calculates for every candidate a Pearson's correlation matrix by default (Spearman and Kendall are also possible to use). At that point, it can have for every targeted protein complex a Pearson's correlation matrix. The raw matrix for each correlation matrix consists of a list that correlates the intensity detected in the mass spectrometer for each fraction with the ID of each protein.

##### 1.2 mCP – Prefilter stage

After that, a prefilter stage is initiated. On the correlation matrixes, a fixed filter is applied. The first filter is based on a minimalistic definition as follows: A protein complex is a minimum composition of at least two proteins that have a significant co-elution hit. A significant co-elution hit means that a binary interaction of proteins within the protein complex will have in the correlation matrix a Pearson correlation coefficient higher than the fixed filter threshold.

When the prefilter stage is finished, mCP plots all detected candidates. These protein complex candidates have at least two components with one significant binary interaction hit (higher than the filter).

##### 1.3 mCP – FDR evaluation

The mCP function provides two possible search modalities. The *de novo* search (dynamic=FALSE) and the dynamic (dynamic=TRUE). Both search modalities are explained in Supplementary Material 1. The dynamic search is intended for the analysis of known and curated PPCs, while the other search is intended for actual *de novo* searches.

Here we will briefly explain the dynamic search. All detected candidates from the pre-filter search are regarded as a valuable source of information about the characteristics of each protein complex. The number and Pearson's correlation average of significant hits for each candidate are annotated. This new threshold is used in the Monte Carlo simulations, where mCP searches are repeatedly performed on the decoy matrixes. Here, the Pearson's coefficient is dynamically adjusted for each candidate according to the intrinsic characteristics of the selected candidates detected in the pre-filter search.

The result of each simulation is a list of all candidates detected in the pre-filter stage that displays how many significant hits does the decoy complex contains. Again, this time the Pearson's correlation threshold filter is specific for each protein complex since the correlation matrix is known for all candidates. The false discovery rate (FDR) is then calculated according to the following equation:

$$FDR=FP/MCs$$

where FP represents the number of detected false positives, and MCs is the number of Monte-Carlo simulations performed. The specific protein complex will be set as a false positive if the number of significant hits in the simulation is equal to or higher than in the experimental candidate complex.

##### 1.4 mCP – Monte-Carlo decoy matrices

Decoy matrixes are generated to detect false positives in each Monte-Carlo simulation. Once the decoy matrices are generated, they will be sequentially run through the whole mCP workflow with a dynamic filter. The first step in the decoy generation is to eliminate all proteins in common between CORUM and the experimental dataset to avoid these known interactions. On the matrix that remains, the IDs intersected between the experiments and the CORUM database will be replaced at random into the experimental matrix. That way, decoys will have a random profile each time.

##### 1.5 mCP – Outputs

Once FDR evaluation is finished, mCP will only include protein complexes that were equal to or lower than the false discovery rate limit assigned. If no FDR threshold is specified, the output will include the protein complex along with it respective FDR evaluation. The following outputs files will be generated:

1) A plot displaying the profiles of all potential protein complexes during the chromatographic separation, resembling Fig. 1 A. On the x-axis, mCP will plot the molecular weight calibrated to the fraction number, depending on the information provided. On the y-axis, the intensities detected by the mass spectrometer will be plotted. If the setting relative = TRUE in the mCP function is added, relative abundance plots will be generated. Normalization plots are achieved within each protein complex by dividing the maximum intensity achieved by each protein into each fraction. Therefore, the resulting plots will have a relative scale ranging from zero to one on the y-axis for each protein of the detected protein complex.
2) a pdf file with plots of heatmaps and network heatmaps for all potential protein complexes (candidates). Plots are similar to the ones shown in Figure 2, in the panel labeled 'Output'.
3) the same plots explained in 1 and 2 for the detected protein complexes after the FDR filter is applied.

4) a *.txt record file with setting parameters used in the analysis.
5) a *.csv register of the FDR Monte Carlo simulation.
6) the main output table (*.csv file) that includes protein complexes detected, significant hits, FDA for each protein complex, and useful parameters described in supplementary information 1.

Note: mCP contains a monomeric filter that takes out potential protein complexes if the maximum of the signal is present in a fraction where the monomers are expected. This is related to the previous filter, which by default is off so the user can decide (details can be found in the R package), running the command: help(mCP).

#### 2 Optimizing settings to maximize complexes detection

Once the wet lab protocol has been performed, there are two major steps carried out for detecting protein complexes. The first is the detection and quantitation of precursors, peptides, and protein groups using established DIA-MS processing softwares (Fig. 1 B-D); the second is the detection of PPCs by the mCP platform (Fig. 1 E). mCP has three main settings (out of a total of 19 parameters) located in the main mCP function that have a significant influence on PPC detection, namely: 1. method_cor, 2. filter and 3. N_simulations.

These settings can be selected in the main function of the R package. We described and provide recommendations for the DIA platform (Fig.1) and the mCP settings shortly here. For more detailed information on the other parameters see Supplementary Material 1.

##### 2.1 Filter settings for mCP

We experimentally determined that a Pearson's correlation coefficient threshold of 0.81 produced the best results. To that end, we performed mCP analysis on our 4 data sets with 185 Monte Carlo simulations each, and varied the filter from 0 to 0.99, as shown in Figures (Suppl. Fig. 11 A and B). The proportion of false positives (or discarded complexes) strongly decreased when the first filter was increased up to 0.81.

In addition, we performed an analysis of the same datasets with a fixed filter of 0.81, i.e. with the dynamic filter turned off, and observed that the filter does not markedly affect the number of detections, but starts taking out protein complexes that will pass the FDR filter at higher values. So, the number of detections decreases continuously when the filter takes numeric values higher than 0.81 (Suppl. Fig. 11B). Therefore, filter = 0.81 takes out potential false positives in the prefilter search and reduces the number of candidates to be evaluated in the FDR simulation.

Other possible and valid options are to set the filter to values of e.g. 0 or 0.5, because three out of the four data sets in this study are derived from sample P2R1. This allowed for a direct comparison between DIA-MS processing software platforms, with performance decreasing in the order 1) DIA-NN, 2) Spectronaut and 3) Max Quant (Fig. 2E, Suppl. Fig. 11, Suppl. Fig. 12).

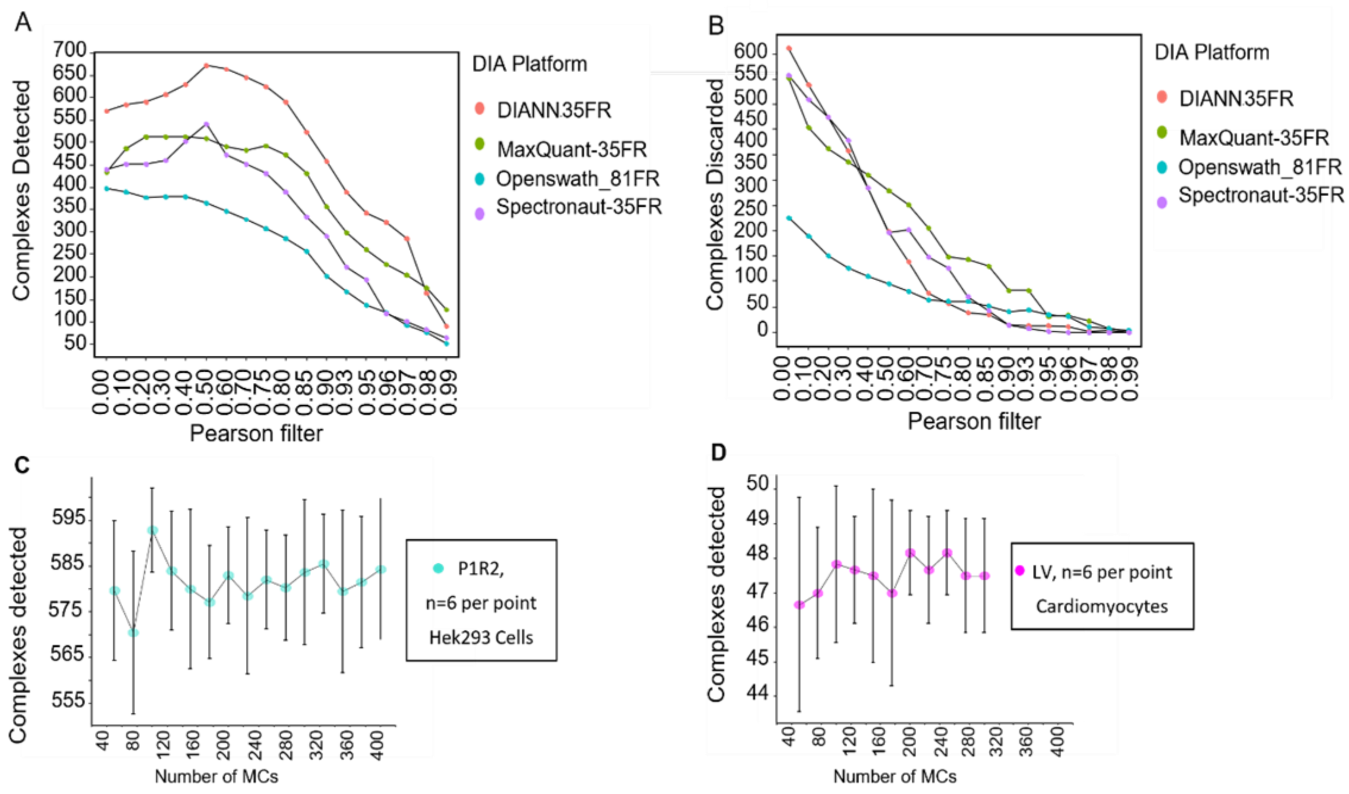


**Supplementary Figure 11: Settings for mCP analysis using the dynamic search modality.** A) mCP sensitivity vs. Pearson's correlation coefficient fixed filter values on different DIA platforms for P1R2 (Fig. 1) and the 81 fraction dataset (Heusel et al., 2019). B) Discarded protein complexes vs. Pearson's correlation coefficient fixed filter values on different DIA platforms C), D) Number of MCs used for FDR assessment for HEK293 and mouse left ventricle cardiomyocyte experiments, respectively.

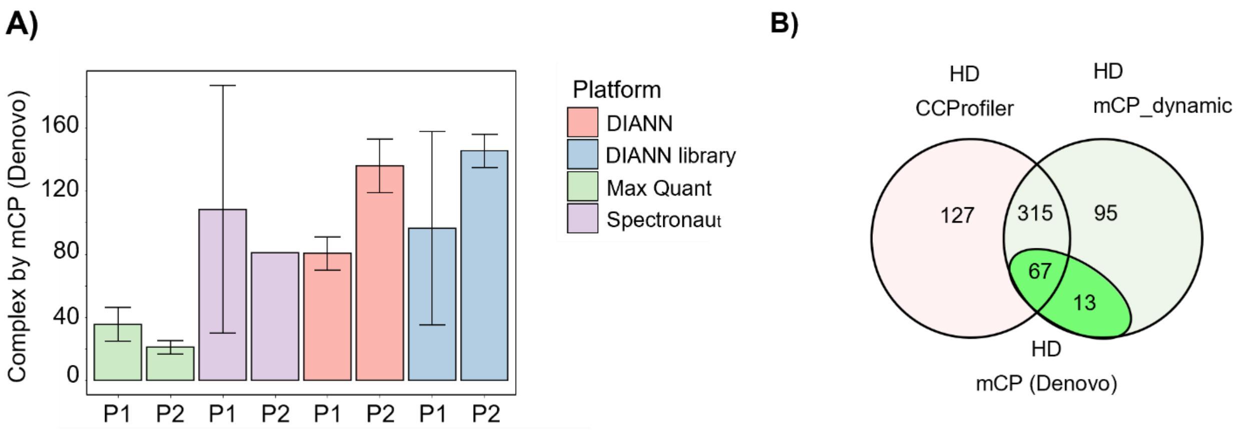


**Supplementary Figure 12: mCP *de novo* search modality.** A) Performance comparison of different DIA-MS processing software packages on the detection of protein groups from Hek293 cells P1 and P2. B) Venn diagram of complexes detected by the mCP R package on an 81 fraction data set (Heusel et al., 2019) by mCP dynamic search vs. mCP *de novo* search vs. CCprofiler search.
